# Galvanin (TMEM154) is an electric-field sensor for directed cell migration

**DOI:** 10.1101/2024.09.23.614580

**Authors:** Nathan M. Belliveau, Matthew J. Footer, Amy Platenkamp, Celeste Rodriguez, Heonsu Kim, Christopher K. Prinz, Aaron P. van Loon, Yubin Lin, Tara E. Eustis, Michelle M. Chan, Daniel J. Cohen, Julie A. Theriot

**Affiliations:** Department of Biology and Howard Hughes Medical Institute, University of Washington, Seattle, WA 98195, USA; Department of Molecular Biology, Princeton University, Princeton, NJ 08544, USA; Department of Electrical and Computer Engineering, Princeton University, Princeton, NJ 08540, USA; Lewis-Sigler Institute for Integrative Genomics, Princeton University, Princeton, NJ 08544, USA; Department of Mechanical and Aerospace Engineering and Omenn-Darling Bioengineering Institute, Princeton University, Princeton, NJ 08540, USA

## Abstract

Directed cell migration of immune and epithelial cells is critical for their rapid response to tissue injury or infection. Endogenous electric fields generated by disruption of the transepithelial potential across the skin have been postulated to play an important role in guiding cells to wound sites, though how individual cells sense these tissue-scale physical cues remains largely unknown. We have identified Galvanin (TMEM154), a previously uncharacterized single-pass transmembrane protein, as being required for electric-field-guided migration of individual rapidly moving cells. Galvanin functions in both immune and epithelial cell types. Upon exposure of cells to an electric field, Galvanin rapidly relocalizes to the anodal side of a cell, and the net charge on its extracellular domain is necessary and sufficient to drive this spatial relocalization. Furthermore, expression of Galvanin is sufficient to confer electric field–guided migration on otherwise non-responsive epithelial cells. In human neutrophils, we show that Galvanin relocalization is immediately followed by changes in the spatial pattern of cellular protrusion and retraction. The strong directional response of these cells is lost upon truncation of Galvanin’s intracellular domain, suggesting that Galvanin acts as a direct sensor of the electric field, transducing spatial information about a cell’s electrical environment to the intracellular migratory apparatus. This sensor relocalization mechanism of cell steering defines a new paradigm for directed cell migration.

## Introduction

Directed cell migration is important for many aspects of biology. In mammals, directed migration drives processes critical to development, immune system function, and tissue regeneration after injury^1–3^. Much of our understanding of how migrating cells effectively sense and respond to environmental signals comes from work on chemotaxis, mediated by transmembrane receptors that bind specific chemical ligands and transduce those signals to reorient the mechanical force-generating components of the cytoskeleton, and thereby direct persistent directional migration ^4,5^. Less is understood about how cells sense and integrate other available spatial information about their physicochemical environment, such as gradients in temperature, pH, or matrix stiffness^6^.

Directed cell migration in response to acute injury presents a particularly interesting challenge, as the injury must generate a new kind of spatial information directing cells to move toward the wound that was not present in the pre-existing normal structure of the tissue. Here, wound-induced electric fields represent one such directional signal. Nearly all polarized animal epithelia maintain distinct ionic environments on their apical versus basolateral sides due to asymmetric ion transport, and the epithelial barrier presents electrical resistance, normally resulting in a transepithelial potential difference^7,8^. Acute disruption of epithelial barrier integrity by physical injury causes a short-circuit in the transepithelial potential, generating a wound-induced endogenous electric field with magnitudes of 50-500 mV/mm that can persist for many hours^9–11^. Many cell types, including epithelial cells and tissue-resident immune cells, have been found to migrate directionally in response to such electrical cues in a process called electrotaxis (or galvanotaxis)^12^, which has been proposed to play an integral role in the wound healing response^13^.

Although it has been known for well over a century that many motile cell types can perform electrotaxis, the molecular mechanism of electric field detection remains ill-defined. To mediate long-distance directed cell migration, a directional signal generated at the cell surface must be transduced to the mechanical elements of the cytoskeleton. Indeed, several intracellular signaling components known to be involved in chemotaxis have also been shown to contribute to electrotaxis^12,14,15^, indicating that electrotaxis is a cell-regulated biological response rather than a purely physical response to forces generated by the influence of electric fields on charged macromolecules. However, electrostatic shielding and poor conductance across the plasma membrane make it unlikely that the relatively weak wound-induced electric fields directly affect the localization or activity of intracellular signaling components^16–19^.

A complicating factor in understanding cellular electric field detection is the diversity in electrotactic behavior across different cell types, electric field magnitudes, and modes of cell movement (individual versus collective)^16,20^. Even within a single cell type, the sensitivity and directional preference in response to an electric field can depend on group size^23–26^.

Madin-Darby Canine Kidney (MDCK) cells, for example, form large epithelial sheets capable of collective cathodal electrotaxis, but these cells respond poorly to electric fields as individual, isolated cells^25^. Recent work in the collective electrotaxis of neural crest cells in *Xenopus laevis* has identified the voltage-sensitive phosphatase, VSP^15^. Loss of VSP disrupts the electric-field-dependent dissemination of neural crest cells in embryos. However, while ex vivo clusters also exhibit electrotaxis, isolated single cells are unresponsive to an electric field, suggesting particular distinctions in the mechanism of collective versus single cell electrotaxis. For individual slowly moving cells, such as fibroblasts, with typical migration speeds of only a few tens of microns per hour, the EGF receptor and several types of ion channels have been implicated in electrotaxis, although it is not clear if any of these act primarily as direct electric field sensors^21,22^.

For very rapidly moving cells capable of responding to electric fields as individual cells (rather than as cell clusters or larger tissues), including immune cells and skin epidermal cells, that crawl approximately two orders of magnitude more swiftly than fibroblasts^16,27^, the identity of a dedicated electric field sensor protein capable of supporting rapid electrotaxis responses has remained elusive. Biophysical experiments and theoretical modeling have most strongly supported a hypothesis that cells can sense the presence and orientation of electric fields via spatial redistribution of charged cell surface macromolecules by electrophoresis and/or electroosmosis in the plane of the plasma membrane^16,28,29^. We therefore sought to discover cell surface proteins that act directly as electric field sensors in such cells. Here, we develop an unbiased functional genomics approach to isolate human neutrophil-like cells based on their electrotaxis behavior and identify new proteins involved in electric field sensing. We uncover a conserved, cell surface-localized electric field sensor that redistributes rapidly in response to electric cues and mediates directed cell migration. This discovery reveals a new molecular mechanism for electric field detection that operates across diverse cell types.

## Results

### CRISPR-interference screen of electrotaxis in neutrophil-like cells identifies Galvanin (TMEM154)

Human neutrophils and neutrophil-like cell lines exhibit impressively rapid amoeboid directional migration in response to a wide variety of chemical cues^30^, and also migrate toward the cathode in an applied electric field^13^. The HL-60 cell line, originally derived from a patient with promyelocytic leukemia, can be differentiated in culture to a neutrophil-like cell phenotype^31^ and is amenable to genome-wide unbiased genetic screens using CRISPR or CRISPR-interference (CRISPRi) approaches^27,32^. We engineered a device to spatially separate millions of differentiated HL-60 cells based on their capacity to migrate toward the cathode upon exposure to physiologically relevant electric field strengths (Fig. 1A, further detailed in Fig. S1-S2). Cells were added to the top of a membrane containing 3 μm diameter pores and directed downward by applying an electric field with the anode above and the cathode below the membrane. We could then collect both the cells that successfully migrated through the membrane from the bottom reservoir and those that remained above. Application of an electric field of about 200 mV/mm substantially increased the fraction of cells that could be recovered in the bottom reservoir, with approximately 60% of the initial cell population recovered after two hours (Fig. 1B). In contrast, undirected migration in the same device in the absence of an electric field resulted in recovery of only 15% of the initial cell population in the lower reservoir.

**Figure 1.**
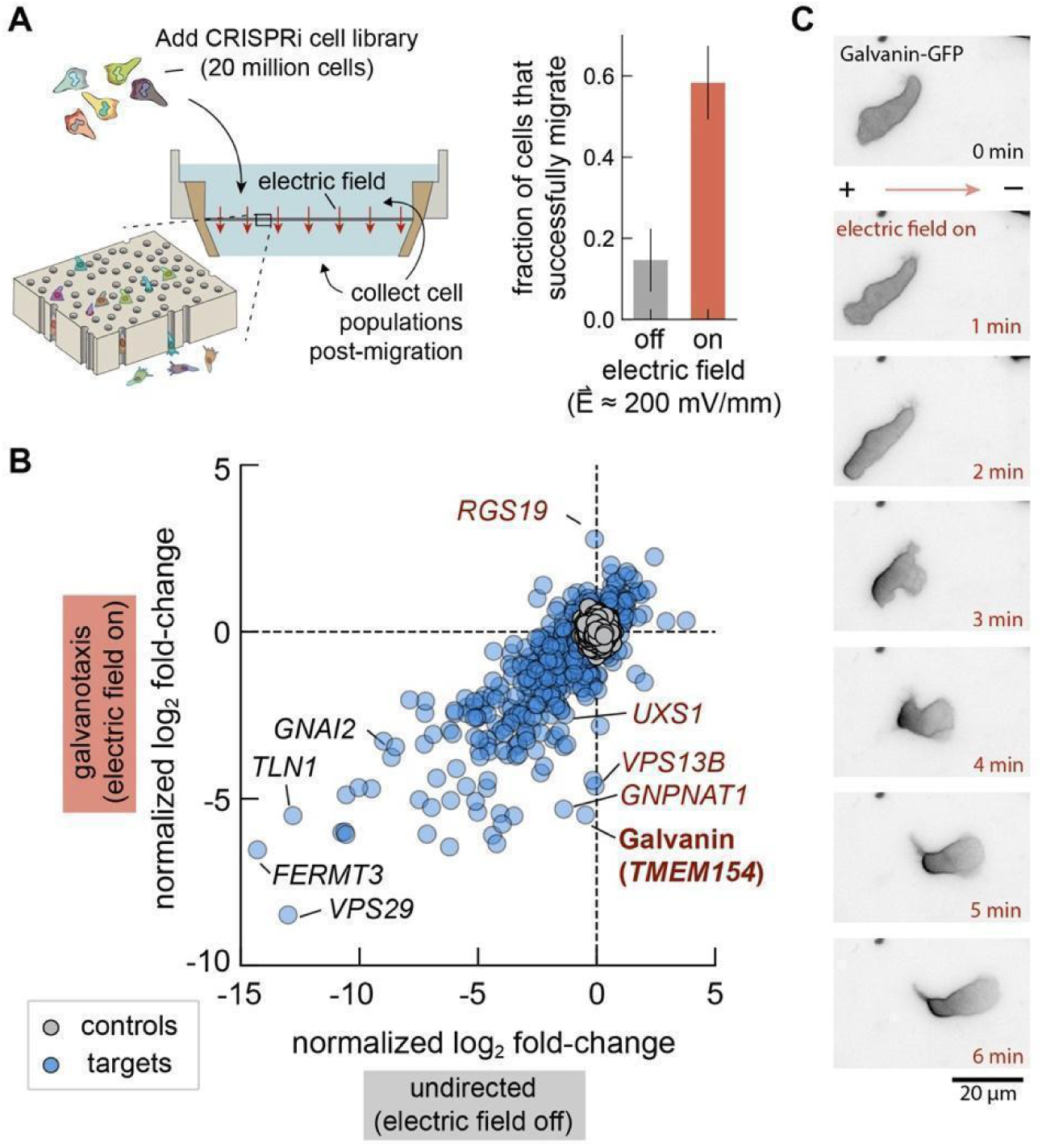
CRISPR interference screen identifies Galvanin as a putative electric field sensor for directed cell migration. **A.** Left: Schematic of galvanotaxis screen. A pooled CRISPRi library of dHL-60 neutrophil-like cells migrate through a membrane with 3 μm diameter pores. Collection of the sub-population that migrate through and, separately, those that remain above the membrane, allow sgRNA target enrichment and gene candidate identification. Right: Fraction of cells collected in the bottom reservoir after two hours (p-value = 0.001, two-sided Mann-Whitney U test). **B.** Summary of focused library screen targeting 1,070 genes for knockdown. Data points show the normalized log_2_ fold-change averaged across three sgRNAs per gene across independent experiments (11 replicates for galvanotaxis and 10 replicates for undirected migration). Control values were generated by randomly selecting groups of three control non-targeting sgRNAs. **C.** Fluorescence micrographs show rapid localization of Galvanin (TMEM154)-GFP toward the anodal pole when differentiated HL-60 neutrophils are exposed to an electric field (300 mV/mm). See also *Supplemental Movie 1*.

We performed a genome-wide unbiased genetic screen in two phases. Initially, we used a genome-wide CRISPRi cell library including sgRNAs targeting 18,901 genes. This was used to narrow our focus in a secondary screen where we targeted 1,070 genes whose knockdown showed the strongest effects on cell migration probability. This approach allowed us to increase the coverage per sgRNA in our secondary screen, and thereby improve our statistical confidence for the effects of individual gene candidates (see *Supplemental Methods*). To better distinguish between perturbations specific to sensing the electric field versus those affecting cell migration more generally, we compared results from two experimental conditions: a) exposure of cells to an electric field (200 mV/mm) and b) undirected cell migration in the same device in the absence of an electric field. In the secondary screen, we found 473 genes where CRISPRi knockdown significantly disrupted migration in the presence of an electric field and 544 genes whose knockdown significantly affected undirected migration, relative to control sgRNAs present in the library (Fig. 1B; adjusted p-value with cutoff of 0.05). The majority of these candidate genes had significant effects in both conditions (Pearson correlation *r* = 0.8 between normalized log_2_ fold-change values), reflecting the commonality of migration machinery necessary during neutrophil migration regardless of the nature of the external cue^27,33,34^. A smaller subset of 111 genes, however, were identified as significantly altering cell migration probability only in the presence of the electric field. This subset included *TMEM154* (transmembrane protein 154), that we refer to as Galvanin, which stood out as the transmembrane protein whose knockdown gave the strongest electric-field-specific phenotype.

Galvanin is predicted to be a single-pass transmembrane protein (161 amino acids, UniProt ID: Q6P9G4). In primary neutrophils, its transcriptional levels are roughly similar to other integral membrane proteins such as the LPS receptor CD14 and the integrin ɑ ^35^. We confirmed that Galvanin localizes to the HL-60 cell plasma membrane by tagging the intracellular C-terminus with eGFP. As a putative electric field-sensitive protein, we hypothesized that its localization would become spatially biased when cells were exposed to an electric field. Using an agarose overlay to confine migrating cells to a single plane on a coverslip, we exposed cells to an electric field of 300 mV/mm. We indeed observed relocalization of Galvanin-GFP, which rapidly becomes biased to the anodal (positive) pole of the cell within about one minute of exposure (Fig. 1C, Supplemental Movie 1). For these cells, which migrate toward the cathode, this equates to relocalization of the protein to the cell rear, and is consistent with electrophoresis of a net negatively charged protein. From the amino acid sequence alone, the ectodomain is expected to have a net charge of-7*e*, which is too low to account for such a notably biased distribution^16^. One likely explanation for this disparity is glycosylation of the ectodomain, which could substantially increase its negative charge. Protein glycosylation, referring to the addition of carbohydrates to the polypeptide following translation, has been implicated in electrotaxis of other cell types^16,28,36^. These modifications often form branched oligosaccharide chains that terminate in negatively charged sialic acid groups and can confer a strongly negative net charge on glycosylated proteins at neutral pH ^37^(*24*). Using NetNGlyc 1.0^38^ and NetOGlyc 4.0^39^, we found six amino acid positions in the ectodomain of Galvanin that may contain glycosylation (one N-type and up to five O-type modifications).

We followed up on several of the screen candidate genes to better assess how the genetic perturbations altered cell migration. In addition to Galvanin itself, we focused on three other candidate genes that might be expected to affect the cell surface presentation of a putative electric field sensor: VPS13B which is involved in membrane trafficking, and GNPNAT1 and UXS1 which are both enzymes involved in protein glycosylation. We generated individual stable HL-60 knockdown lines for each of these four candidates, then differentiated them and embedded them in three-dimensional collagen gels, where their speeds and migration directions were measured using video microscopy. Consistent with our screen results, each cell line appeared to migrate normally in the absence of an electric field. Importantly, in the presence of an electric field, each cell line showed reduced directionality when compared to migration of cells with a non-targeting sgRNA (Fig. S2E), supporting their involvement in the directional response.

### Galvanin is important for the directional migration response during electrotaxis of human neutrophil-like cells

To better assess the functional contribution of Galvanin to directional migration, we used CRISPR-Cas9 to create two clonal knockout HL-60 cell lines where the endogenous Galvanin gene was completely disrupted (Fig. S3A). We then assessed their directional migration in collagen gels when exposed to electric fields of varying strength. In Fig. 2A we show tracks of individual migrating cells exposed to a 300 mV/mm field, tracking cell nuclei over a 30-minute period. We observed a notable loss of directed migration in each Galvanin knockout cell line relative to wild-type cells (Fig. 2Ai middle, Fig. S3B). In Fig. 2Aii, we replot these cell tracks with a common starting origin. Importantly, by expressing our Galvanin-GFP construct in the Galvanin knockout cell line, we were able to rescue the cathodal directional migration of wild-type cells (Fig. 2Ai and Fig. 2Aii, right), demonstrating that Galvanin is necessary for the normal electric field response. Notably, the Galvanin knockout cells were still highly migratory, with average migration speeds indistinguishable from wild-type cells in the presence or absence of an electric field (Fig. 2B); that is, their phenotypic defect lies in their ability to orient toward the cathode and not in any other aspect of cell migration. We further quantified the average speed of individual cells projected along the electric field vector (that is, the electric field-directed component of the cell speed) (Fig. 2C). Consistent with the differences observed in the cell tracks, we find a significant loss of directed movement in our knockout cell line compared to the wild-type and Galvanin rescue cell lines.

**Figure 2.**
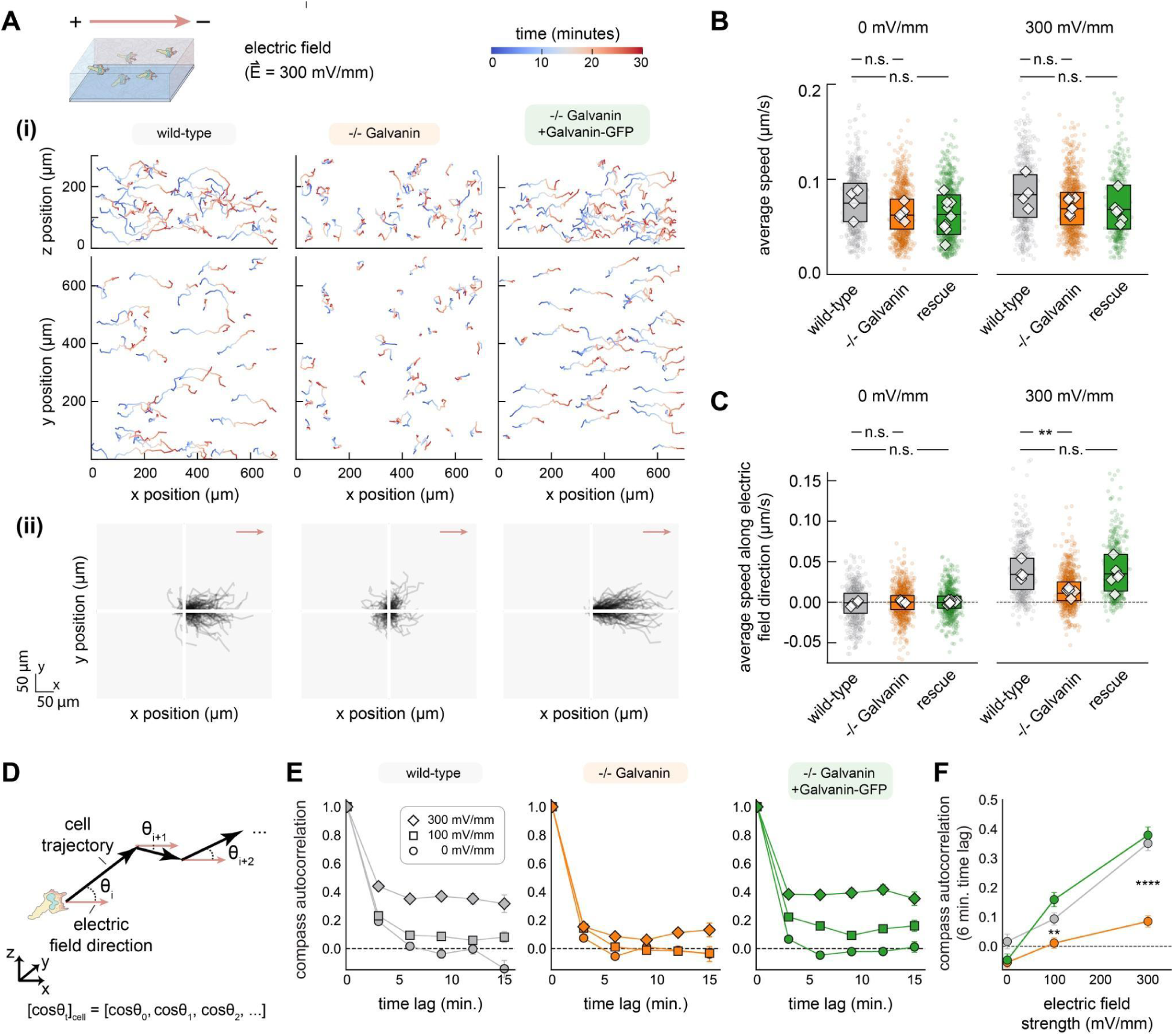
Galvanin is critical for persistent cathodal migration in HL-60 neutrophils. **A.** Nuclear tracking was performed during migration of HL-60 neutrophils in a collagen gel exposed to an electric field (300 mV/mm). (i) Data represents a subset of data, with only 50 cell tracks shown per cell line (dHL-60 neutrophil wild-type,-/- Galvanin knockout, and the genetic rescue with Galvanin-GFP). (ii) The same tracks are replotted starting at 0,0. Red arrow indicates the direction of the electric field. **B.** Average speed calculated based on nuclear tracking with 3 minute time intervals (two-sided Mann-Whitney U test found no significant difference across experimental replications). **C.** Average speed parallel to the electric field direction calculated based on nuclear tracking with 3 minute time intervals. A significant difference is noted between the Galvanin knockout and wild-type cells when exposed to a 300 mV/mm field (p-value=0.006, two-sided Mann-Whitney U test). For **B** and **C,** Individual colored scatter points represent average values of individual cells, while the boxplot extends from the first to third quartiles with a line at the median. The white diamonds represent average values across cells from experimental replicates). **D.** Schematic showing calculation of compass autocorrelation, used to assess longer-term directed movement. **E.** Compass autocorrelation. For individual cell tracks, the angle 𝜃 between the cell vector and the electric field vector was determined for each 3 minute time interval trajectory. Autocorrelation analysis was performed on the set of corresponding cosine 𝜃 values and averaged across cells. Error bars represent standard error of the mean. **F.** Autocorrelation values for a six minute time lag. Error bars represent standard error of the mean. A significant difference is noted between the Galvanin knockout and wild-type cells (100 mV/mm: p-value=0.003, 300 mV/mm: p-value<0.0001, two-sided Mann-Whitney U tests comparing individual cell values). For each cell type in **B**, **C**, **E**, and **F,** 400-1,000 cells were analyzed per condition (approximately 30-150 cells quantified per imaging acquisition, with 4-9 acquisitions per cell line and per electric field strength).

We also assessed the directional movement at longer length scales. For migration in a physically complex environment such as fibrous collagen gel, which results in somewhat tortuous cell trajectories, we chose a metric to assess the overall directional bias under a sustained cue. Specifically, we quantified the autocorrelation of measured cosine of the angles (cosine 𝜃) between the migration vector of individual cells and the electric field vector, which we refer to as the compass autocorrelation (Fig. S2D). This metric remains high when cells move over long periods of time in the same net direction, even if the local path persistence is relatively low (as is typical for rapidly moving neutrophils). Cells exhibited a compass autocorrelation that was dependent on the strength of the electric field and was notably reduced with the Galvanin knockdown cell lines (Fig. SE). In Fig. 2F we plot the compass autocorrelation at a six-minute time lag, with the Galvanin knockout cell line showing a substantial loss of directional movement along the electric field vector, and complete rescue by Galvanin-GFP. We conclude that Galvanin is an indispensable element for directed migration during electrotaxis of these rapidly moving neutrophil-like cells.

### Galvanin contributes to electrotaxis in other metazoan cell types

We next considered whether Galvanin is functional in other cell types. Transcriptomics data suggests that Galvanin is predominantly expressed in immune cells and epidermal tissue^40,41^. Among immune cells, Galvanin appears most highly expressed in granulocytes, though it is also observed in other cell types including macrophages, T cells, and B cells. Within epidermal tissues, basal and suprabasal keratinocytes also show high levels of expression. Such an expression profile is consistent with a putative role in processes related to skin injury and wound healing.

We first turned to T cells, another immune cell type that performs amoeboid migration and that has been found capable of electrotaxis^42^. Here we generated a clonal Galvanin knockout in the highly motile *Mus musculus* EL4 T cell line^43^ using CRISPR-Cas9 in the same manner as our HL-60 neutrophils above. To assess their response to an electric field, we assessed migration in collagen gels using video microscopy as for the HL-60 neutrophils. Individual cell tracks are plotted, with a common origin, for wild-type and Galvanin knockout EL4 T cells exposed to a 300 mV/mm field (Fig. 3A). Similar to the neutrophils, we find that these cells exhibit a cathodal directional preference, with a notable defect in cathodal movement following Galvanin knockout.

**Figure 3.**
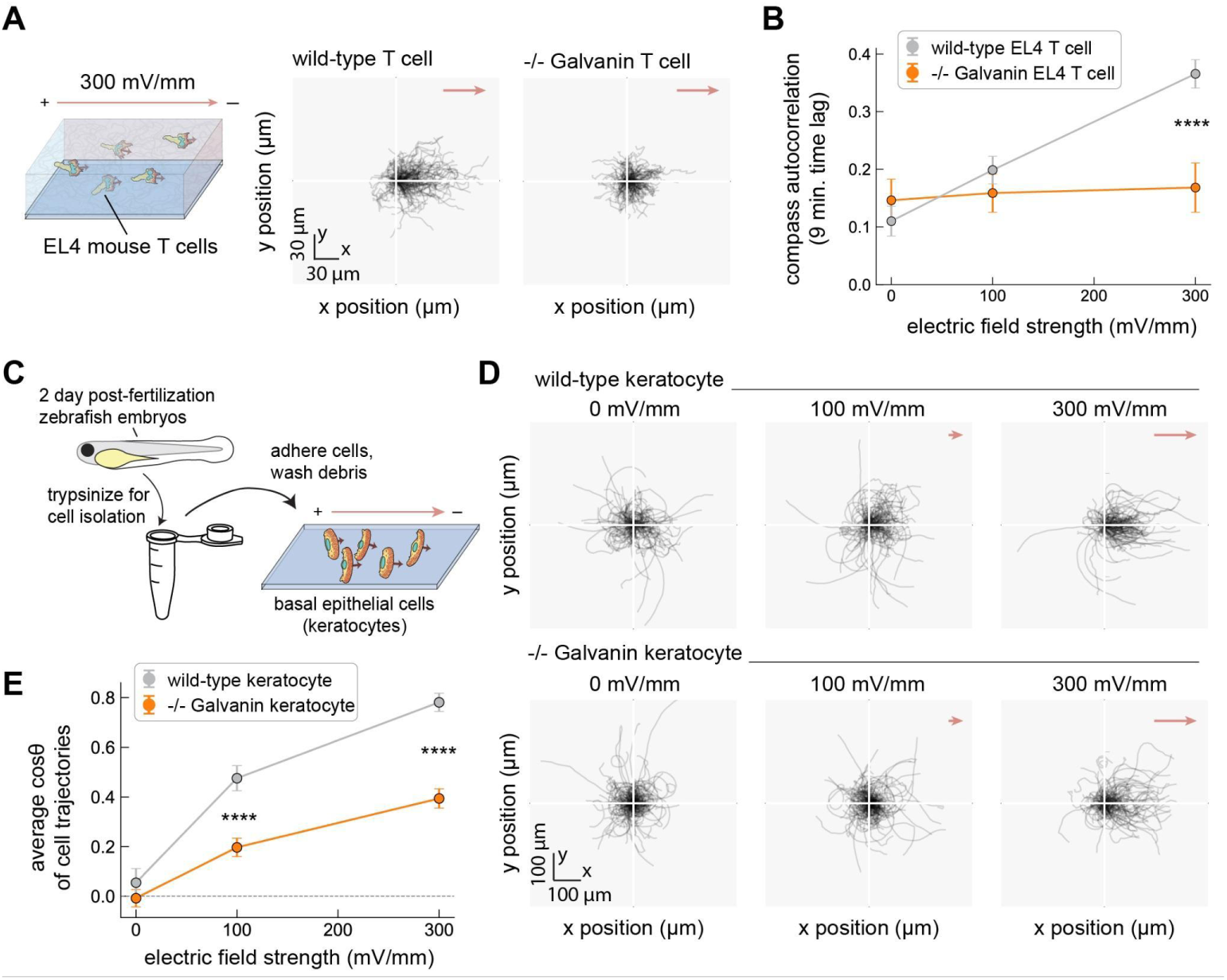
Galvanin contributes to electrotaxis in other cell types and other vertebrate animal species. **A.** Left: schematic showing murine EL4 T cells embedded in a collagen gel. Right: Relative cell trajectories of wild-type and Galvanin knockout EL4 T cells exposed to a 300 mV/mm field (60 minutes tracking, 2 minute imaging interval). 200 cell trajectories are shown per condition, which represent a random subset of the data for visual clarity. **B.** Compass autocorrelation of cells migrating in a collagen gel, associated with data shown in part **A** (9 minute time lag). 295-956 cells per cell line and per electric field condition (N= 6 sample preparations for wild-type, and 3 sample preparations for-/- Galvanin). Error bars represent standard deviation of the mean. Asterisks indicate statistical significance between the individual cell measurements from wild-type and-/- Galvanin EL4 cells (p < 0.0001, two-sided Mann–Whitney U test). **C.** Schematic showing isolation of migratory fish keratocytes. Keratocytes were isolated from 2 day post-fertilization zebrafish embryos (wild-type or-/- Galvanin) and allowed to adhere to coverslips prior to electrotaxis experiments. **D.** Relative cell trajectories of keratocytes exposed to an electric field (300 mV/mm) over 30 minutes (1 minute imaging interval). A subset of trajectories are shown (125 per condition). **E.** Average cosine of the angle 𝜃 between cell trajectories (final and initial position) and the electric field vector. Error bars represent standard error of the mean. Analysis was performed across 126-373 cell trajectories per condition, from 4 sample preparations. Asterisks indicate statistical significance between the individual cell measurements from wild-type and-/- Galvanin keratocytes (p < 0.0001, two-sided Mann–Whitney U test).

Using our compass autocorrelation metric, we find a significant decrease in biased movement toward the cathode in the Galvanin knockout cell line versus wild-type EL4 T cells (Fig. 3B). This result demonstrates that Galvanin contributes to cathodal electrotaxis in other mammalian immune cell types.

We also wanted to assess whether Galvanin contributed to electrotaxis in epidermal tissue and turned our attention to the basal epidermal cells from *Danio rerio* (zebrafish). These cells, often referred to as fish keratocytes, have been used extensively as a model system to study rapid single-cell electrotaxis and exhibit a cathodal directional preference^16,44^. Galvanin knockout fish were generated by CRISPR-Cas9 excision of a large portion of the Galvanin gene locus in single-cell embryos (Fig. S4). In order to assess keratocyte electrotaxis, we isolated primary cells from homozygous knockout embryos two-days after fertilization and seeded them onto coverslips (Fig. 3C). Fig. 3D shows individual cell trajectories from wild-type and Galvanin knockout cells across 0 mV/mm, 100 mV/mm, and 300 mV/mm field strengths. We observe a reduced directional bias, with greater cell track density apparent around the origin of Galvanin knockout keratocytes exposed to 100 mV/mm and 300 mV/mm fields, though residual cathodal directionality remains apparent. To quantify the directional response during 2D migration, it is common to calculate the average cosine 𝜃 across a population of cells, where 𝜃 represents the angle that a cell travels relative to the electric field direction. Here, a value of 1 indicates a complete directional response toward the cathode, while a value of 0 indicates no response on average. In Fig. 3E, we plot the average cosine 𝜃, finding a significant reduction in the cathodal bias of Galvanin knockout cells over wild-type cells. In summary, we find that Galvanin is functional in a variety of cell types relevant to skin injury and wound healing. Furthermore, by exploring its function for cell types in mouse and zebrafish, we show that Galvanin contributes to electrotaxis more broadly among metazoan species.

### Expression of Galvanin is sufficient to enable cathodal migration of single cells in an electric field

Our results so far demonstrate that expression of Galvanin is *necessary* for the strong cathodal directional bias observed in the cell types considered so far. We were next interested in testing whether Galvanin is *sufficient* to mediate cathodal bias of cells exposed to an electric field. Here we turned to Madin-Darby Canine Kidney (MDCK) cells. While these cells exhibit cathodal electrotaxis during collective migration as an epithelial sheet, they have been found to respond poorly as single cells^25^. These cells express little to no detectable Galvanin^45^. We hypothesized that if we expressed Galvanin in these cells, the protein would exhibit anode-directed membrane redistribution following exposure of cells to an electric field, and then aimed to determine whether single cells might become more responsive to electric fields.

To test these hypotheses, we engineered two MDCK cell lines expressing increasing amounts of human Galvanin-GFP, as measured by flow cytometry (Fig. 4A, see Methods). To assess Galvanin localization and migratory behavior, cells were sparsely seeded on a cover slip as single cells or small cell clusters (approximately 2-10 cells). We began by looking at the expression and spatial redistribution of Galvanin-GFP by live-cell video microscopy. Consistent with our observations in HL-60 neutrophils, we observed Galvanin localization to the plasma membrane, and a notable redistribution of Galvanin toward the anodal pole across both isolated single cells and cell clusters under application of an exogenous electric field (Fig. 4B). To quantify migratory behavior, we tracked cell nuclei during 2D migration upon exposure of cells to a field strength of 300 mV/mm for three hours. Consistent with prior work and counter to the cathodal movement observed in with larger epithelial sheets of MDCK cells^25^, the wild-type individual cells and small clusters exhibit poor electrotaxis, though we note an apparent modest anodal bias across the population (Fig. 4C, top). Strikingly, expression of Galvanin-GFP resulted in a notable cathodal switch in migration direction (Fig. 4C, middle and bottom).

**Figure 4.**
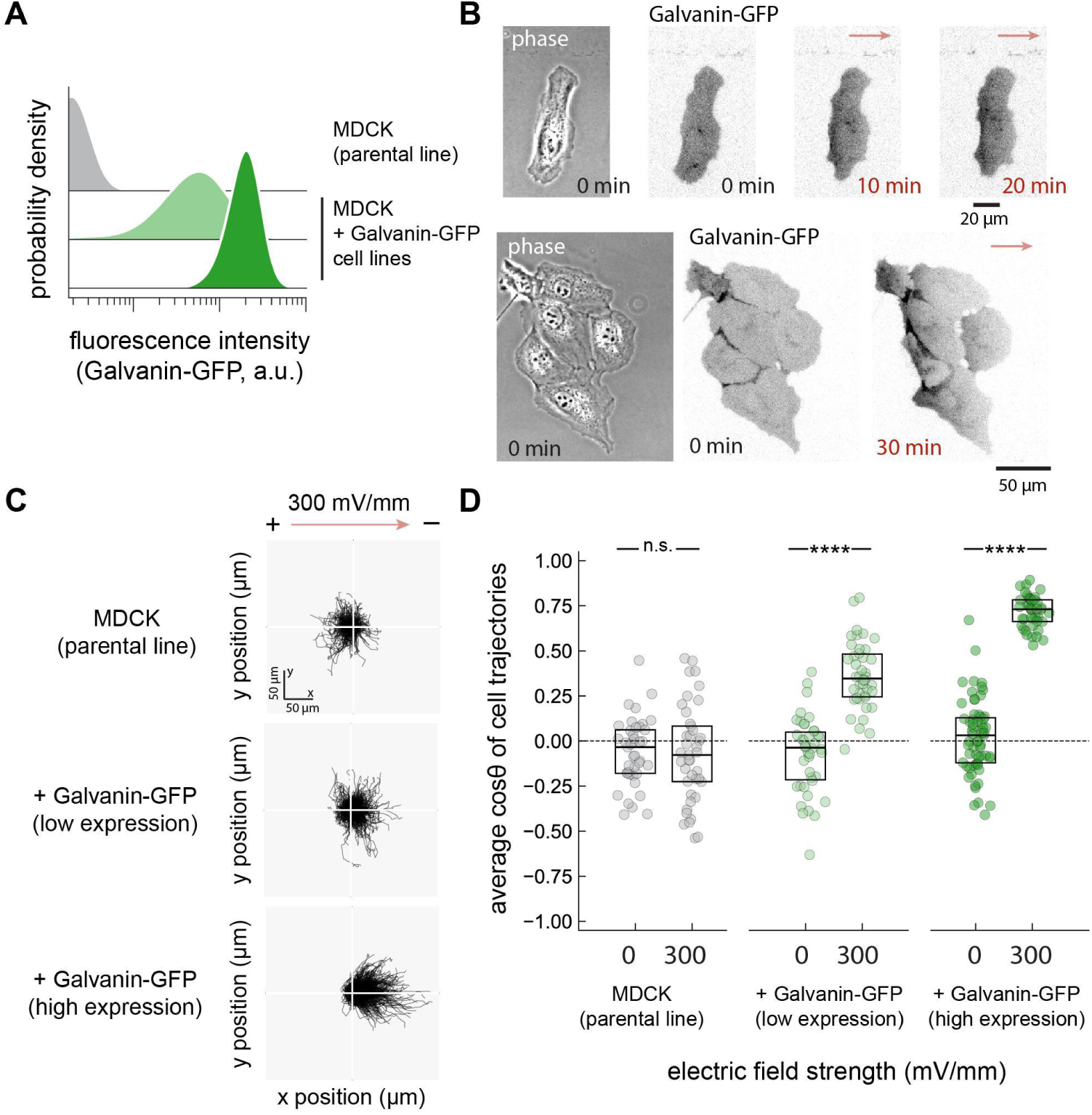
Galvanin is sufficient to enable cathodal electrotaxis. **A.** Flow cytometry histograms showing expression of human Galvanin-GFP in two clonal MDCK cell lines. The parental MDCK cell line is not fluorescent and their intensity values reflect autofluorescence (see Methods). **B.** Fluorescence micrographs showing localization of Galvanin-GFP toward the anode when MDCK cells are exposed to an electric field (300 mV/mm). *left*: example of a single cell. *right*: example of a cell cluster. **C.** Relative cell trajectories of MDCK cells exposed to an electric field (300 mV/mm) over three hours (5-minute imaging interval). A subset of trajectories is shown (100 per condition). **D.** Average speed along the electric field direction. Each data point represents the speed averaged across cells within a single imaging field of view (40 fields of view across two sample preparations; 971-1760 cells per condition), while the boxplot extends from the first to third quartiles with a line at the median. Asterisks indicate statistical significance between the 300 mV/mm and 0 mV/m conditions (p-value < 0.0001, two-sided Mann–Whitney U test).

Quantifying the directional bias by the average cosine 𝜃 across cell trajectories, as with the fish keratocytes above, we observed a significant Galvanin dose-dependent increase in the cathodal bias of MDCK cells (Fig. 4D). We also separately quantified the behavior of single cells and small cell clusters, observing a similar Galvanin dose-dependence on cathodal migration (Fig. S5). Interestingly, the movement of single cells from the parental line exhibited a significant anodal bias during exposure to a 300 mV/mm field, which was strongly reversed toward the cathode with increasing Galvanin expression. This shows that, in addition to supporting cathodal electrotaxis, Galvanin is able to override alternative electrotaxis mechanisms. In sum, our results demonstrate that Galvanin is both necessary and sufficient to drive cathodal electrotaxis for a wide variety of motile cell types across multiple vertebrate animal species.

### Relocalization of Galvanin on the plasma membrane defines the cell front and cell rear during electrotaxis

To better understand how Galvanin supports directed migration, we returned to our HL-60 neutrophils to better quantify the dynamics of its relocalization on the membrane while also measuring changes in local protrusion and retraction activity and front-rear cell polarization^46,47^. Using our under-agarose assay to confine cells to migrate in a single plane, we monitored cell migration at high spatial and temporal resolution during five minutes of undirected migration, five minutes of exposure to an electrical stimulus of 300 mV/mm, and five minutes of recovery (Fig. 5A). Galvanin-GFP localization was measured around the periphery of individual cells, and cell migration changes were quantified using the Adapt package^48^, which measures the relative local protrusion or retraction of the cell edge, as a function of position around the cell perimeter. This analysis was performed on 46 cells, enabling us to generate population-averaged kymographs of Galvanin localization dynamics (Fig. 5B) and an associated mapping of protrusion and retraction (Fig. 5C).

**Figure 5.**
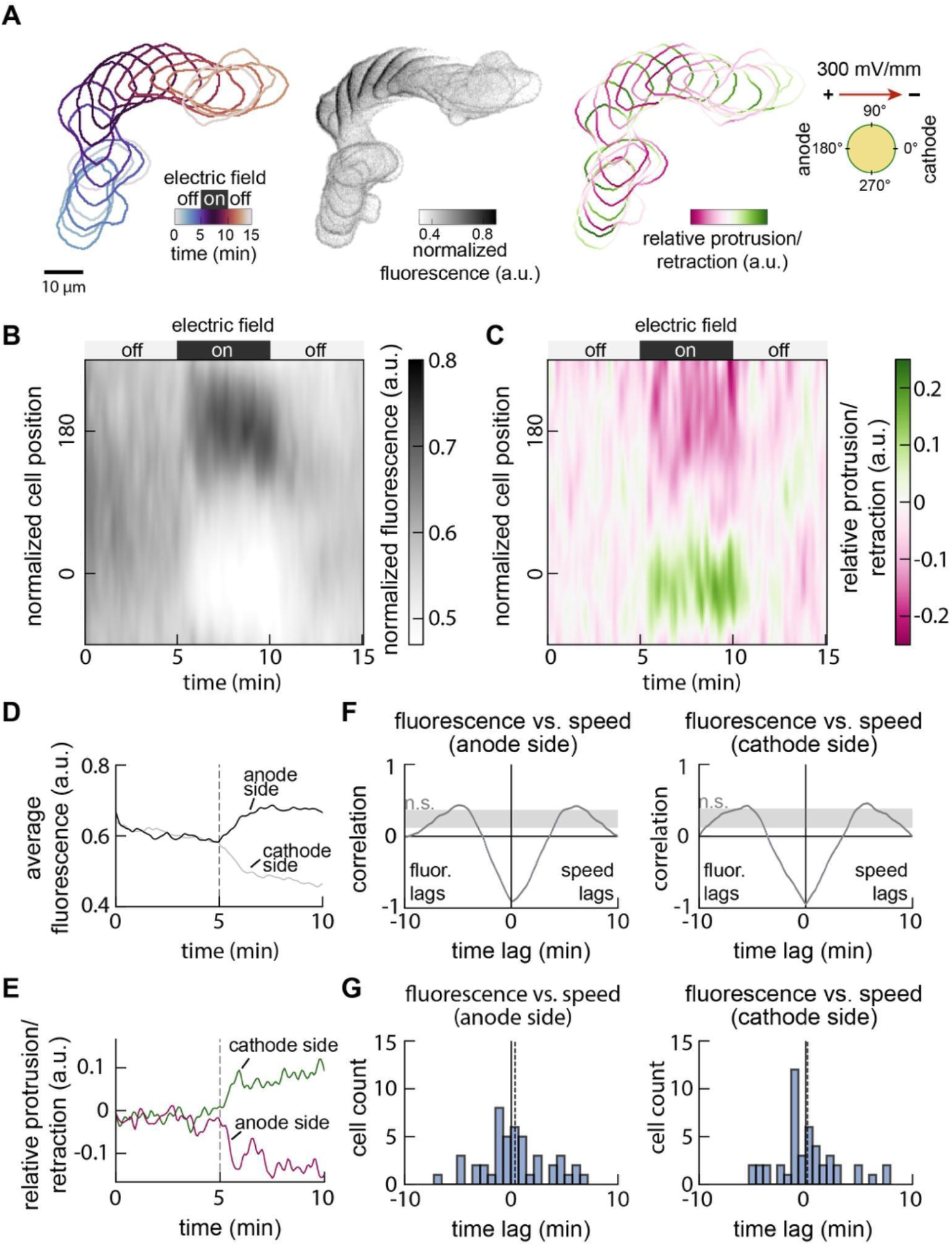
Localization of Galvanin coincides with reduced local membrane speed and front-rear polarization. **A.** Example analysis showing the temporal movement, Galvanin-GFP fluorescence intensity, and cellular protrusion/retraction activity of a single cell. Imaging was performed every 5 seconds over 15 minutes, with cells exposed to an electric field (300 mV/mm) between 5 minutes and 10 minutes. One-minute intervals are overlaid. See also *Supplemental Movie 2*. **B.** Kymograph shows the normalized Galvanin-GFP intensity, quantified at the cell periphery (in a band of 2 μm thickness) and averaged across 41 cells. Cell contours are defined with 360 positions, with position 0 corresponding to the right-most position relative to the cell centroid. **C.** Kymograph quantifies membrane speed along the cell periphery, by comparing cell shape changes between 5 second intervals and correcting against bulk cell translocation. Values represent an average across the 41 cells considered in part B. **D.** Averaged fluorescence at the anodal side (positions +120 to +240) and cathodal side (positions-60 to +60) during the first 10 minutes. **E.** Averaged membrane speed at the anodal side (positions +120 to +240) and cathodal side (positions-60 to +60) during the first 10 minutes. Dashed lines in **D** and **E** indicate when the electric field was turned on. **F.** Cross-correlation analyses comparing the averaged fluorescence and membrane speeds (parts D-E) at the anodal and cathodal sides of the cell. Dashed lines indicate the average minimum lag (0.1 minutes for the anode side and 0 minutes for the cathode side). The shaded region represents correlation values that are below a 99% confidence interval (see *Methods*). **G.** Cross-correlation analysis was performed using the Galvanin-GFP fluorescence and membrane velocities averaged across all 46 individual cells. Histograms show the lag corresponding to the minimum correlation value. Dashed lines near zero indicate the average lag value (anode side: 0.4 minutes; cathode side: 0.2 minutes).

We found that the cellular response to the electric field is almost immediate, with a relocalization of Galvanin-GFP to the anode that was nearly complete within about one minute (Fig. 5D).

Accompanying this was a notable change in the spatial distribution of protrusion and retraction, with an immediate increase in retraction at the newly forming cell rear and an increase in protrusion at what becomes the cell front (Fig. 5E). To compare the timing of the Galvanin-GFP relocalization with the changes in protrusion and retraction activity, we performed a cross-correlation to compare these two parameters over time (Fig. 5F). On both the rear (anodal) and front (cathodal) sides of the cell, the Galvanin-GFP relocalization was tightly coupled to the protrusion/retraction activity, with a maximum negative correlation at zero time lag, indicating that the two processes are nearly simultaneous (within the time resolution of these experiments). This finding was also maintained when measuring the cross-correlation between Galvanin-GFP signal and protrusion/retraction activity for individual cells (Fig. 5G). These results suggest that Galvanin relocalization defines the cell front and cell rear for directed cell migration during exposure to an electrical cue, either by locally activating retraction at the rear, or by removing inhibition of protrusion at the front.

### Relocalization of Galvanin depends specifically on the net ectodomain charge

In theory, the rapid anode-directed electrophoresis of Galvanin should depend on the Coulombic interaction between the charged ectodomain of the protein and the electric field, with the field unable to penetrate into the intracellular space^18,49,50^ (Fig. 6A). The stable spatial distribution of Galvanin-GFP we observed after several minutes of electric field exposure should represent a balance between the Coulombic movement toward the positive, anodal pole with electrophoretic velocity *v_E_* and equilibration by diffusion with an effective diffusion coefficient D^16^. Specifically, we expect the steady-state Galvanin concentration to vary along the electric field vector with an approximate exponential decay^51^, with a characteristic length that depends on the ratio *v_E_*/D. This ratio can be used to infer the net charge on the protein (see *Supplemental Methods*).

**Figure 6.**
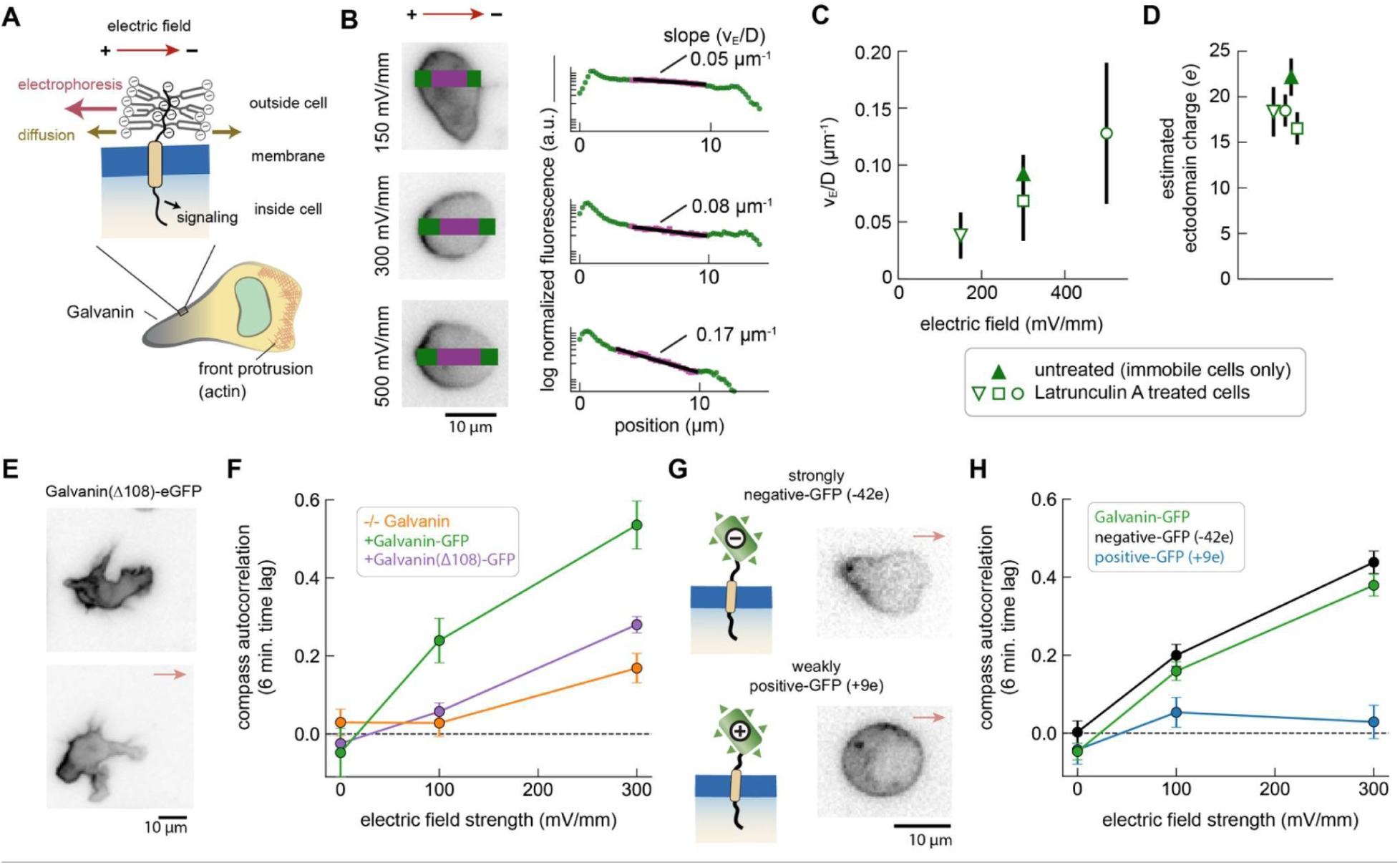
The sensory mechanism of Galvanin depends on a highly charged ectodomain and its intracellular domain. **A.** Schematic illustrating our model of Galvanin electrophoresis due to the net negative charge. Branched changes represent expected glycosylation of ectodomain based on 6 predicted O-/N-type modifications. **B.** Left: Example micrographs of Galvanin-GFP in individual HL-60 neutrophils cells exposed to an electric field for 10 minutes. Cells were treated with Latrunculin A to immobilize them following initial migration under agarose. The intensity profile along the electric field vector direction was quantified to estimate the slope in the decay in fluorescence intensity, expected to be proportional to v_E_/D. Right: semilogarithmic plots of corresponding fluorescence profiles for the examples shown, with green indicating the entire cell width, and magenta corresponding to the fit region, selected to avoid the more intense cell periphery. The estimated v_E_/D values are indicated for each fit. **C.** Summary of v_E_/D values across the different electric field strengths. Error bars indicate standard deviation across 13 cells (150 mV/mm), 23 cells (300 mV/mm), and 26 cells (500 mV/mm). Analysis of non-motile cells present in the experiments of Figure 3 are also included (4 cells). **D.** Estimated ectodomain charge based on the v_E_/D values of part B (see *Methods*). Error bars indicate standard error of the mean. **E.** Micrographs of Galvanin with an endodomain truncation (Galvanin(Δ108)-GFP) showing localization to the plasma membrane, with anodal bias following 3 minutes exposure to a 300 mV/mm electric field. **F.** Compass autocorrelation of cells migrating in a collagen gel (-/- Galvanin: orange;-/- Galvanin + Galvanin-GFP (green),-/- Galvanin + Galvanin(Δ108)-GFP. Data points represent average across analysis performed on individual cell tracks (80-685 cells per cell line and per electric field strength); error bars: standard deviation of the mean. **G.** Left: the ectodomain was altered using engineered GFP proteins with either a strongly negative charge (-42*e* net ectodomain) or weak charge (+9*e* net ectodomain charge). right: Representative fluorescence micrographs of cells expressing the engineered Galvanin constructs exposed to a 300 mV/mm electric field (-42*e,* n=40 cells; +9*e*, n=31 cells). The strongly negative construct shows localization to the anodal side of the cell, similar to the wild-type Galvanin-GFP, while the weakly positive construct remains uniformly distributed. **H.** Compass autocorrelation of Galvanin knockout HL-60 neutrophils expressing engineered constructs: wild-type Galvanin-GFP (green),-42GFP-Galvanin (black), or +9GFP-Galvanin (blue). Data points represent average across analysis performed on individual cell tracks (180-750 cells per cell line and per electric field strength); error bars: standard deviation of the mean.

To more precisely quantify this ratio and avoid confounding factors introduced by migrating cells (including possible membrane flow), we repeated the under-agarose experiments in the presence of Latrunculin A. This drug inhibits filamentous actin assembly and allows us to immobilize cells as they begin to migrate under the agarose overlay, maintaining their flattened shape against the coverslip. Latrunculin A-treated cells were exposed to electric field strengths of 150 mV/mm, 300 mV/mm, and 500 mV/mm (Fig. 6B). The slope of the fluorescence intensity profiles, plotted on a semilogarithmic scale, were then used to estimate the ratio *v_E_*/D at each electric field strength. As shown in Fig. 6C, the *v_E_*/D ratio increased from about 0.05 μm^-1^ to 0.2 μm^-1^ over this range. The net charge is proportional to the *v_E_*/D ratio, normalized by the electric field strength, and strikingly, we arrive at a similar estimate of the net charge on Galvanin across all conditions, with a value of-18*e* (±1.1 SEM) averaged across all the data (Fig. 6D). Using this same dataset, we were also able to estimate the effective diffusion coefficient D for Galvanin-GFP by quantifying its equilibration back to a uniform distribution after the electric field stimulus was removed (Fig. S6A-D), obtaining a value for D of 0.53 μm^2^/s (±0.1 SEM), which is consistent with expectations for a single-pass transmembrane protein^52^. In summary, Galvanin’s high mobility and high net negative charge enable it to exhibit rapid dynamics as an electric field sensor that mediates electrotaxis.

Our results above suggest that asymmetrical redistribution of Galvanin regulates actomyosin activity, allowing cells to navigate toward the cathode. While the signal transduction pathway downstream of Galvanin relocalization remains unknown, it is likely that the intracellular domain plays an important role, either through interaction with cytoskeletal regulatory proteins, or through direct interaction with actin and myosin. To test this hypothesis directly, we generated a construct truncating the entire intracellular domain, Galvanin(Δ108)-GFP and expressed this in our HL-60 Galvanin knockout cell line. We find that this protein exhibits similar anodal redistribution as the full-length protein upon exposure of cells to an electric field (Fig. 6E).

However, when quantifying directional bias in our collagen migration assay, we find that this construct is unable to rescue the directional response of wild-type HL-60 neutrophils (Fig. 6F). These results suggest that, following spatial redistribution of Galvanin, its intracellular domain indeed plays a critical role in regulating the cell’s subsequent directed movement response.

Due to the limited structural characterization of Galvanin, it remained unclear whether additional features of the protein might be important for its sensory function, or whether the net charge of the ectodomain was by itself sufficient. To more definitively assess this, we removed the wild-type ectodomain and replaced it with engineered protein domains that have a known net negative charge (Fig. 6G). Specifically, we utilized previously developed “super-charged” GFP proteins^53^. We hypothesized that a highly negatively charged engineered ectodomain would rescue directed migration in our Galvanin knockout cell line, assuming net charge is the sole critical factor. We were able to express a highly negative construct, replacing Galvanin’s ectodomain with a negatively charged XTEN linker domain^54^ and negatively charged version of GFP, yielding an expected total net charge of-42*e* at pH 7. Additionally, we expressed a construct where the ectodomain was replaced with a weakly positive GFP, having an expected net charge of +9*e* at pH 7. Although these mutated forms of GFP exhibited poorer fluorescence, with the +9*e* form showing more variable membrane and cytosolic fluorescence, both constructs still localized to the plasma membrane in Galvanin knockout cells (Fig. 6G).

We assessed the functional activity of these engineered constructs by expressing them in Galvanin knockout cells and exposing the cells to an electric field. Using Latrunculin A as noted above to immobilize cells, the-42*e* construct showed a strong anodal localization (Fig. 6G, top). Estimating v_E_/D as described above, we calculate a net charge to be 37e (±2.5 SEM), which is close to the calculated value and gives us confidence in the charge measurement technique. In contrast, the +9*e* construct showed no notable change in GFP distribution when cells were exposed to an electric field (Fig. 6G, bottom), suggesting that the weaker charge is insufficient for relocalization at these electric field strengths, as had been previously predicted based on theoretical considerations^16^. To assess migration in cell lines expressing these engineered constructs, we turned to our cell tracking assay in collagen. Importantly, the-42*e* construct was able to completely rescue directed cell migration during exposure to an electric field when compared to cells expressing the wild-type Galvanin construct (Fig. 6H). In contrast, and consistent with the lack of spatial relocalization of the +9*e* construct, the weakly charged ectodomain was unable to rescue the directed migration response. In summary, we find that the high net negative charge of the ectodomain is sufficient to mediate the sensory response of Galvanin to support its functional role in electrotaxis.

## Discussion

We have identified Galvanin, a previously poorly characterized membrane protein, that acts as an electric field sensor and mediates cathode-directed cell migration. We demonstrate that loss of Galvanin expression disrupts directed migration in response to an electric field. In amoeboid-like neutrophil-like cells and T cells, we assessed this during migration in a three-dimensional collagen environment, representative of their migration in the interstitial space of tissue. For the more adherent fish keratocytes and MDCK epithelial cells, we tracked cells as they migrated on a two-dimensional surface. While past work has demonstrated that several plasma membrane proteins undergo electrophoresis across cells exposed to an electric field ^16,28,36,55–57^, it has been challenging to connect bulk protein electrophoresis to electrotaxis or to directed migration at the whole-cell level^58^. Our demonstration that electrophoretic redistribution of a single protein based on the net charge of the ectodomain is necessary and sufficient to reorient directed cell migration represents the first description of an electric field sensor of this class.

Galvanin’s electrophoretic response is rapid, enabling migrating cells to quickly repolarize within minutes of exposure to an electric field. For the rapidly migrating neutrophil-like cell type considered in much of our work, this time scale is consistent with the rapid response at sites of tissue injury. In this context, the wound-induced electric field would provide an additional layer of directional information, alongside other well-characterized chemical guidance cues involved in wound repair^14^.

We show that Galvanin accumulates at the anodal side of individual cells, which becomes the cell rear as cells move toward the cathode. In HL-60 neutrophils, the close coupling we observe between Galvanin relocalization and changes in local cell protrusion and retraction behavior suggests that Galvanin may steer cell migration either by locally activating retraction (for example, by stimulating myosin II contractility) or by locally inhibiting protrusion (for example, by inhibiting activity of the Arp2/3 complex responsible for branched actin network growth). In canonical neutrophil chemotaxis, chemoattractant receptors (typically GPCRs) are uniformly distributed on the cell’s plasma membrane, and ligand occupancy of the receptors is higher on the side of the cell closer to the source of the chemoattractant because of its concentration gradient^30,59^. Downstream signaling from the activated GPCRs leads both to stimulation of actin network assembly at the presumptive cell front and to stimulation of myosin II-based contractility at the cell rear^60,61^. Because of the strong positive feedback within the respective “frontness” and “backness” mechanochemical modules, and strong negative feedback between the modules^62–64^, the cell’s internal signaling architecture promotes robust polarization. In addition, tension in the plasma membrane enforces tight and near-immediate coupling of protrusion at one end of the neutrophil with retraction at the opposite side and vice versa^65,66^. As a result, local activation of retraction or local inhibition of protrusion by Galvanin could each be completely sufficient to reorient whole-cell polarity and drive directed neutrophil migration.

Future work will be needed to understand how Galvanin interacts with and regulates actomyosin activity and whether signaling pathways are shared across the many cell types capable of electrotaxis^13,16,67–69^.

Considering our findings more broadly, accumulating evidence supports the hypothesis that multiple distinct and possibly non-exclusive mechanisms mediate electrotaxis across different cell types and cellular contexts. The identification of the voltage-sensitive phosphatase, VSP, as a key mediator of electrotaxis during directed migration of collective neural crest cells in *X. laevis*^15^, identifies an important putative sensor protein involved in vertebrate development.

However, neural crest cells were only able to perform electrotaxis during collective cell migration; individual cells were non-responsive in applied electric fields. Incidentally, we find only low levels of VSP expression in human neutrophils^35^. A number of additional studies further highlight differences in electrotaxis behavior of solitary motile cells like immune cells versus collective cell electrotaxis as observed for neural crest cells and MDCK monolayers^24–26^.

Collective cell electrotaxis is likely influenced by additional mechanical aspects associated with cell-cell and cell-substrate interactions^23^. Our results in the MDCK cell line provide a striking illustration of gain-of-function in electrotactic responsiveness, where ectopic expression of Galvanin is sufficient to make isolated, individual MDCK cells capable of cathodal electrotaxis. In future work it will be exciting to understand how Galvanin alters the cytoskeletal machinery in single MDCK cells to enforce a switch from anodal to cathodal electrotaxis. It will also be important to understand whether these cells share the same signaling mechanism as rapidly moving immune cells.

In humans, transcriptional data suggests Galvanin is most highly expressed in immune cells and skin, in line with the perceived importance of electrotaxis during wound healing^14^. While further work will be needed to identify the molecular details that allow Galvanin to regulate cellular signaling and cytoskeleton activity, our data demonstrates a biophysical mechanism of protein electrophoresis on the cell surface through direct interaction of the protein’s ectodomain with an external electric field. The ability for cells to alter Galvanin’s net charge through glycosylation, which tends to be cell-type specific^39,70^, also offers an interesting strategy cells can employ to vary its functional activity across different cell types. In the work performed here, our ability to engineer Galvanin and alter its physical and electrical properties demonstrates a tunable biological system that can be used to provide control over directed cell movement.

## Acknowledgements

We thank members of the Theriot lab for useful discussions throughout this work, and we thank Alex Mogilner, Sean Collins and Griffin Chure for comments on the manuscript. We also thank Alexander Leydon and Jennifer Nemhauser for access to their Bioruptor instrument, Takato Imaizumi for access to Bio-Rad equipment used to perform Western blots, and Tom Daniel for access to COMSOL. Research support is provided by the Howard Hughes Medical Institute (J.A.T.), and grants from the National Institutes of Health (N.M.B.: R00GM147355; D.J.C.: R35133547-06). N.M.B. was supported as a Fellow of the Jane Coffin Childs Memorial Fund for Medical Research during some of this work, while D.J.C. and M.C. received additional support from a Schmidt Transformative Technologies Fund.

This article is subject to HHMI’s Open Access to Publications policy. HHMI lab heads have previously granted a nonexclusive CC BY 4.0 license to the public and a sublicensable license to HHMI in their research articles. Pursuant to those licenses, the author-accepted manuscript of this article can be made freely available under a CC BY 4.0 license immediately upon publication.

## Author Contributions

Conceptualization: N.M.B., J.A.T.

Methodology: N.M.B., M.J.F., A.P.

Investigation: N.M.B., T.E.E, A.P., H.K., C.R., Y.L.

Writing - original draft: N.M.B., J.A.T.

Writing - review and editing: M.J.F., A.P., N.M.B., J.A.T.

Supervision: J.A.T., N.M.B., D.J.C., M.M.C.

Funding acquisition: J.A.T., N.M.B., D.J.C., M.M.C.

## Data and Materials Availability

Sequence data generated from CRISPRi screens and RNA-seq are uploaded to the Sequence Read Archive and can be accessed under BioProject accession code PRJNA1147270 (https://www.ncbi.nlm.nih.gov/bioproject/PRJNA1147270). All data files used to generate figures are available in our GitHub repository, noted in the Code Availability section below. Due to the large size of raw image files used for cell tracking, these are not included but are available from the corresponding authors upon request. Source data are provided with this paper. Processed data, code, and figure generation scripts generated with the Python package matplotlib (v. 3.5.2) are publicly available as a GitHub repository (nbellive/CRISPRi_galvanotaxis_pub).

## Supplementary Materials

Materials and Methods

Figs. S1 to S6

Movies S1 to S2

Data S1

