## Supplemental Information for "Galvanin (TMEM154) is an electric-field sensor for directed cell migration"

**The PDF file includes:**

Materials and Methods
Figs. S1 to S6
Captions for Movies S1-S2 and Data S1

Other Materials for this manuscript include Supplemental Movies S1-S2 and Supplemental Data S1.

### Table of Contents

**Materials and Methods** **3**
Cell Culture and Zebrafish husbandry 3
Cloning and Cell Line Generation 3
Pooled Plasmid Library Synthesis 3
Cloning of individual sgRNA plasmids for CRISPRi 5
Lentivirus Production for Stable Integration of sgRNA 5
Generation of CRISPRi Cell Lines 6
Generation of Galvanin Knockout Cell Line via CRISPR-Cas9 6
Generation of Galvanin Knockout Zebrafish via CRISPR-Cas9 8
Genotyping of Galvanin Knockout in Zebrafish 9
Generation of Galvanin-eGFP expressing Cell Lines 10
Genome-Wide and Secondary CRISPRi Assays 11
Development of Device for Pooled Screens 11
Overview of Cell Collection and Experimental Replicates 12
Removal of Cell Debris and Dead Cells Prior to Cell Migration Assays 13
Cell Migration Screen Assay 13
Quantification of sgRNA from CRISPRi libraries and Gene Identification 13
Image Acquisition 14
Three-Dimensional Migration in Collagen Extracellular Matrices 15
Experimental 15
Cell Tracking and Quantification 16
Two-Dimensional Migration 17
Zebrafish Keratocytes: Cell Isolation, Slide Preparation, and imaging 17
MDCK epithelial cells: Slide Preparation and Imaging 17
HL-60 Neutrophils: Slide Preparation and Imaging 18
HL-60 Neutrophils: Quantification of Peripheral Galvanin-GFP Signal and Membrane Velocity 18
HL-60 Neutrophils: Quantification of vE/D Ratio and Estimation of Charge 19
HL-60 Neutrophils: Quantification of Galvanin's Diffusion Coefficient 21
Immunolabeling for Western Blots 22
Antibodies and Dilution Information 22
**Supplemental Figures and Figure Captions** **23**
Figure Supplement 1 23
Figure Supplement 2 25
Figure Supplement 3 27
Figure Supplement 6 30
**Captions for Supplemental Items** **31**
Supplemental Movie 1 31
Supplemental Movie 2 31
Supplemental Data Table 1 31

### Materials and Methods

#### Cell Culture and Zebrafish husbandry

Undifferentiated HL-60 cells were a gift from the Weiner lab, University of California, San Francisco. EL4 cells were obtained from the American Type Culture Collection (ATCC, #TIB-39). Wild-type MDCK-II cells were originally obtained from the Nelson Laboratory, Stanford University.

HL-60 and EL4 cells were cultured in RPMI 1640 medium containing L-glutamine and 25 mM HEPES 1640 (Gibco #22400089) supplemented with 10% heat-inactivated fetal bovine serum (hiFBS, Gemini Bio Products #900–108), 100 U/mL penicillin, 10 µg/mL streptomycin, and 0.25 µg/mL Amphotericin B (antibiotic-antimycotic, Gibco #15240). MDCK cells were cultured in low-glucose DMEM (Gibco, 1 g/L glucose, #11885084), 100 U/mL penicillin, 10 µg/mL streptomycin (Gibco, 15140122), and 10% fetal bovine serum (FBS, Gemini Bio Products #900–108). All cells were maintained at 37℃ in 5% $CO_2$ in humidified air.

Differentiated HL-60 cells were generated by incubating cells in media containing 1.57% dimethyl sulfoxide (DMSO, Sigma, #D2650). Here, approximately 1 - 1.5 x $10^6$ cells/mL were diluted by adding two volumes of additional media containing the DMSO. This culture media was replenished with fresh media, including DMSO, three days following the initiation of differentiation, and used in cell migration assays five days following the initiation of differentiation.

The transgenic TgBAC(ΔNp63:Gal4, UAS:LifeAct-EGFP) zebrafish line[71,72] was used for this work, with fish maintained according to standard procedures[73]. Experiments were approved by University of Washington Institutional Animal Care and Use Committee (protocol 4427-01). Animals were raised on a 12 hr light, 10 hr dark cycle at 28.5°C. Adults were crossed through natural spawning. Embryos were collected and raised in 100 mm petri dishes at 28.5°C. Each dish contained 30–50 embryos in system water.

#### Cloning and Cell Line Generation

##### Pooled Plasmid Library Synthesis

All genomic integrations for the CRISPR interference (CRISPRi) work involved lentiviral transduction using undifferentiated HL-60 cells expressing dCas9-KRAB (pHR-UCOE-Ef1a-dCas9-HA-2xNLS-XTEN80-KRAB-P2A-Bls) as previously described (*18*). dCas9-KRAB is linked by a proteolysis-resistant 80 amino acid XTEN linker, driven by an EF1α promoter that was placed downstream of a minimal-ubiquitous chromatin opening (UCOE) element to prevent gene silencing and is based on a construct originally gifted by Dr. Marco Jost and Dr. Jonathan Weissman.

The genome-wide sgRNA library was reported in Sanson et al. (*55*) (Dolcetto CRISPRi library set A, Addgene #92385). This library contains 57,050 sgRNA, with 3 sgRNA per gene target and 500 non-targeting control sgRNA. For optimal library design, sgRNA were selected based on their position relative to annotated transcription start sites, expected on-target activity, and the presence of off-target matches.

The focused sgRNA library was constructed by taking the 1,070 most significant gene candidates identified in the initial screen. These were selected based on their adjusted p-value using the Benjamini–Hochberg procedure, with an arbitrary significant threshold set to allow selection of the desired number of genes. The sgRNA library was designed using a similar approach as the Dolcetto CRISPRi library, with 3 sgRNA per gene and then another 250 control sgRNA included from the original library. When selected individual sgRNA for gene targets included, those that appeared ineffective (i.e. individual $\log_2$ fold-change values similar to the control sgRNA) were not included. In their place, an alternative sgRNA was selected from the Dolcetto CRISPRi library set B, which was also reported in Sanson et al. (*55*). During filtering, genes specifically associated with transcription, translation and oxidative phosphorylation were also excluded. In addition, sgRNAs targeting the following genes were appended to the library: RICTOR, TRAF6, IFNAR2, PKN1, PRKCB, IL4R, HCAR2, B3GAT3, CHSY1, CHST9, CHST12.

The sgRNA library was synthesized as previously described (*55*). Briefly, oligonucleotides were synthesized to contain the sgRNA sequences, flanking BsmBI recognition sites, as well as primer sites for the initial amplification of the oligonucleotide library: 5′-[forward primer] CGTCTCACACCG [sgRNA, 20 nt] GTTTCGAGACG [reverse primer], where the forward primer and reverse primer sequences are AGGCACTTGCTCGTACGACG (5′ – 3′) and ATGTGGGCCCGGCACCTTAA (5′ – 3′), respectively (Twist Biosciences, CA). The oligonucleotide library was resuspended in 10 mM Tris-Cl, pH 8.5 to a concentration of 10 ng/µL. A 25 µL polymerase chain reaction (PCR) reaction was prepared, with 5 ng library DNA combined with 12.5 µL 2x NEBNext Ultra II Q5 Master Mix (#M0544S, New England Biolabs) and 0.5 µM of the forward and reverse primers noted above (Integrated DNA Technologies, IA). Library amplification was performed as follows: Step 1) Initial denaturation at 98℃ for 30 sec., Step 2) 10 cycles of denaturation at 98℃ for 10 sec.) and anneal/extension at 65℃ for 30 sec., and Step 3) a final extension at 65℃ for 5 min. Amplified DNA was purified using a QIAquick PCR purification kit (#28104, Qiagen) following protocol directions.

Next, the oligonucleotides were ligated into pXPR_050 (Addgene #96925) using a Golden Gate reaction (*56*). The reaction was performed In 10 µL, containing 1 µL T7 10x StickTogether DNA Ligase Buffer, 0.75 µL BsmBI (10 U/µL, #FERER0451, ThermoFisher Scientific), 0.5 µL T7 DNA ligase (#M0318S, New England Biosciences), 135 ng pXPR_050 DNA, 2.7 ng oligonucleotide insert DNA, and bovine serum albumin diluted to a final concentration of 100 µg/mL. This reaction was performed on a thermocycler: Step 1) 15 cycles alternating between 37℃ for 1.5 min. and then 25℃ for 3 min., Step 2) 50℃ for 5 min., and Step 3) 80℃ for 10 min. The DNA was purified using the QIAquick PCR purification kit and resuspended in 25 µL water. 5 µL of the ligation product was electroporated into 20 µL ElectroMAX Stbl4 electrocompetent cells (#11635018, ThermoFisher Scientific). After a 90 minute outgrowth in SOC outgrowth media

(#B9020S, New England Biosciences), cells were plated onto Bioassay LB agar plates (#12-565-224, ThermoFisher Scientific) with 100 µg/mL carbenicillin. This was repeated six times total and colonies from the six plates were scraped to combine. Plasmid DNA was prepared using a plasmid maxiprep kit (performed by Genewiz from Azenta Life Sciences, NJ). Aliquots from each transformation were also used to estimate electroporation efficiency by plating out diluted aliquots onto 10 cm LB agar petri dishes containing 100 µg/mL carbenicillin. A total of 8.1 million unique transformants was estimated.

#### Cloning of individual sgRNA plasmids for CRISPRi

Individual sgRNA plasmids were constructed using sgRNA identified from the genome-wide CRISPRi screens as previously described (*18*). Briefly, pXPR_050 was first linearized using the restriction enzyme BsmBI (New England Biolabs, #R0739S, which includes NEB buffer 3.1). Here, 20 µg of pXPR_050, 20 µL NEB buffer 3.1, and 10 µL BsmBI were combined for a 200 µL reaction and incubated for 5 hours at 55℃. The resulting linear pXPR_050 DNA was gel extracted using the QIAquick gel extraction kit (Qiagen, #28704) and resuspended in TE buffer (10 mM Tris·Cl, pH 8.0; 1 mM EDTA) to a concentration of 10 ng/µL. The sgRNA inserts were generated by annealing complementary oligonucleotides with DNA overhangs compatible for ligation with the BsmBI-digested pXPR_050 DNA. The oligonucleotides were purchased as described below and annealed by combining 1.5 µL of each forward and reverse oligonucleotide (stock concentration of 50 µM in water), 5 µL NEB buffer 3.1, and 42 µL water. The mixture was first incubated for 5 minutes at 95℃ and then allowed to cool by lowering the temperature by 5℃ every 5 minutes until the sample was at room temperature. Finally to insert the sgRNA into the pXPR_050 vector, 1 µL of annealed sgRNA was combined with 20 ng of the BsmBI-digested pXPR_050 DNA and ligated using T4 ligase (New England Biolabs, #M0202S). The ligated DNA product was transformed into NEB Stable Competent E. coli (New England Biolabs, #C3040H) following manufacturer directions and successfully inserted sgRNA were confirmed by Sanger sequencing (performed by Genewiz from Azenta Life Sciences, NJ).

Forward oligonucleotides were 5' CACCG(20 bp sgRNA target sequence)3', while reverse complement oligonucleotides were 5' AAAC(20 bp reverse complement sgRNA target sequence)C 3' (Integrated DNA Technologies, IA). The sgRNA target sequences used for individual CRISPRI knockdown cell lines are listed below.

Control sgRNA: AGGGCACCCGGTTCATACGCNGG
TMEM154 (Galvanin) sgRNA: GGGAACGAGCGCGATCACCA
UXS1 sgRNA: GGTAGGGCCCTGGACCCGCG
GNPNAT1 sgRNA: CGCGCCACAGTTGGGAACCG
VPS13B sgRNA: CCAGGGACTTGGAGGTGGAG

#### Lentivirus Production for Stable Integration of sgRNA

For large scale lentivirus production (dCas9 construct or pooled sgRNA library), 15 µg transfer plasmid, 18.5 µg psPAX2 (Addgene #12260), and 1.85 µg pMD2.G (Addgene #12259) were diluted in 3.5 ml Opti-MEM I reduced-serum media (Gibco #31985070) and then combined with

109 µL TransIT-Lenti Transfection Reagent (Mirus, MIR6600). Following a 10 minute incubation, this mixture was added dropwise to confluent HEK-293T cells (ATCC, CRL-3216) in a T175 flask containing 35 mL DMEM media (Gibco #11965-092) and supplemented with 1mM sodium pyruvate (Gibco #11360-070). Lentivirus was recovered by collecting media 48 hr later, by centrifugation at 500 g for 10 minutes to remove any residual cells and debris. For our dCas9-KRAB construct, we additionally concentrated the lentivirus approximately 60-fold using Lenti-X Concentrator (Takara Bio Inc., #631231).

For small scale lentivirus production (individual sgRNAs), lentivirus was prepared in 6-well tissue culture plates. Here 1 µg sgRNA transfer plasmid, 1 µg psPAX2, and 0.1 µg pMD2.G were diluted in 200 µL Opti-MEM I reduced-serum media and combined with 6 µL transIT. Following a 10 minute incubation, this mixture was added dropwise to confluent HEK-293T cells, with lentivirus collected as noted above.

#### Generation of CRISPRi Cell Lines

The dCa9-KRAB and sgRNAs constructs were integrated into undifferentiated HL-60 cells using a lentivirus spinoculation protocol. Briefly, lentivirus was added to 1 mL cells ($1x10^6$ cells /mL) and polybrene reagent (final concentration of 1 µg/mL) in 24-well tissue culture plates. Cells were spun at 1,000 g for two hours at 33℃. The supernatant was removed and cells were placed in an incubator for two days prior to antibiotic selection for 6 days (dCas9-KRAB: blasticidin 10 µg/mL; sgRNA constructs: puromycin 1 µg/mL).

For CRISPRi sgRNA library preparations, lentiviral titers were estimated by titrating lentivirus over a range of volumes (0 µL, 75 µL, 150 µL, 300 µL, 500 µL, and 800 µL) with $1x10^6$ cells in a total of 1 mL per well of a 24-well tissue culture plate, using the spinoculation protocol noted above. Two days post-transduction, cells were split into two groups, with one placed under puromycin selection. After 5 days, cells were counted for viability. A viral dose that led to a 12.5% transduction efficiency was used for subsequent pooled library work. This low efficiency was targeted to ensure most cells only received one sgRNA integration.

#### Generation of Galvanin Knockout Cell Line via CRISPR-Cas9

Galvanin expression was disrupted by targeting exon 1 (containing the start codon and signal sequence) using CRISPR-Cas9, with clonal cell lines isolated to obtain cells with non-synonymous mutations. This was performed following a similar strategy to prior work in HL-60 cells (*57*) and protocol recommendations using the Alt-R CRISPR-Cas9 system (Integrated DNA Technologies, IA). Transfection was performed using the Neon Transfection System (Thermo Fisher, #MPK5000) using their 10 µL kit (Thermo Fisher, #MPK1096) as described below.

Briefly, the gene targeting crRNA (GGGAACGAGCGCGATCACCA, human HL60; TGGGGCTCGCGTTTGGTCAA, mouse EL4 T cell) was synthesized by Integrated DNA

Technologies. The sgRNA was prepared by resuspending the crRNA and fluorescently tagged tracrRNA (Alt-R™ CRISPR-Cas9 tracrRNA, ATTO 488, 20 nmol; #10007810) in IDTE buffer (IDT, #11-04-02-01), each to a 200 μM concentration. These two RNA were annealed by diluting 2.2 μL of each crRNA and tracrRNA to a total volume of 10 μL in IDTE buffer and heating the sample to 98°C for five minutes and allowing it to cool to room temperature on the bench top. The sgRNA-Cas9 ribonucleoprotein complex was then formed by first diluting 0.3 μL Alt-R Cas9 S.p. Cas9 Nuclease V3 (62 μM stock, IDT, #1081059) with 0.2 μL Resuspension Buffer R (from Neon 10 μL kit). This was combined with 0.5 μL of the annealed RNA sample and incubated at room temperature for 15 minutes. To prepare the final electroporation mixture, 1 μL of RNP complex was combined with $5x10^5$ wild-type HL-60 cells resuspended in 9 μL Buffer R and 2 μL of 10.8 uM Alt-R Cas9 Electroporation Enhancer (IDT, #1075916). Electroporation was performed with the Neon Transfection System according to manufacturer directions (electroporation settings: 1350 V, 35 ms pulse, 1 pulse). A non-sgRNA control was also included. Cells were then placed in 500 μL RPMI culture media containing 10% hiFBS for recovery. Transfection of the RNP was checked approximately 48 hours later by flow cytometry and confirmed by an observed increase in fluorescence from the fluorescently tagged tracRNA.

Gene disruption was initially confirmed with the polyclonal cells through Sanger sequencing using the TIDE protocol (*58*), which estimates the distribution based on the read peak intensities, sgRNA target site, and known wild-type sequence. This suggested a high prevalence of single and double base-pair deletions (approximately 70% for HL-60 cells and 80% for EL4 cells). Clonal isolates were then generated by limiting dilution (HL-60 cells) or single cell sorting (EL4 cells) into 96 well tissue culture plates and growth in RPMI media containing 50% hiFBS.

Genotyping was determined by Sanger sequencing. For human HL-60 cells, the annotated exon 1 for TMEM154 (Ensemble, ENST00000304385) is ACGTTTCAGAGAGGCTGCAGCCCGGCGCAGCATCCTGAGCGCGCCTCTGCCGAGGCGAGCGGAC**ATGATGCAGGCTCCCCGCGCAGCCCTAGTCTTCGCCCTGGTGATCGCGCTCGTTCCCGTCGGCCGGG**, where the bolded text indicates the coding sequence. Clone 1, used in the work of the main text, was found to have one allele with single base-pair deletion in the coding sequence (**…CCTGG-GATCGCGCTCGTTCCCGTCGGCCGGG**), where the dash indicates the deletion. The other allele contained two larger deletions, with the first spanning the starting codon (ACGTTTCAGAGAGGCTGCAGCCCGGCGCAGCATCCT GAGCGCGCC- - - - - - - - - - - - - - - - - - - **- - - - - - - - - - - - - - - - - - - - - - - CTAGTCTTCGCC- - - - - - - - - - - - -CTCGTTCCCGTCGGCCGGG).** Clone 2, a secondary cell line used to confirm loss of directionality in an electric field (Fig. S3A) contained the same single base-pair deletion as clone 1, but at both alleles (**…CCTGG-GATCGCGCTCGTTCCCGTCGGCCGGG**). For mouse EL4 cells, the annotated exon 1 for TMEM154 (Ensemble, ENSMUSG00000056498) is AGTCTGACAGGCTCCTGCAGAGTTACATCACTTCCTAAAAATCAGGCAAACACCTCTCCTTCAAGTGAGCACCGACCCAAAGGAGCAGGAAACAGGAAATGTGCGCGTTGGAGGGAGGCCGGGGCTTCGCAGCATCCTGAGCGCAGGCGAGCCAGCTGAC**ATGACCGTGCCCTGTGCTGCGTTGGTCTTGGCCCTGGGGCTCGCGTTTGGTCAATCCAGCCAGG**, where the bolded text indicates the coding sequence. The isolated clone had alleles with either a single base-pair or two base-pair deletion in the coding region(**ATGACCGTGCCCTGTGCTGCGTTGGTCTTGGCCCT-GGGCTCGCGTTTGGTCAATCCAGCCAGG and ATGACCGTGCCCTGTGCTGCGTTGGTCTTGGCCCT- -GGCTCGCGTTTGGTCAATCCAGCCAGG)**, where the dash indicates the deletion.

#### Generation of Galvanin Knockout Zebrafish via CRISPR-Cas9

The zebrafish Galvanin knockout fish was generated by excising a portion of genomic DNA spanning the gene's transcription start site up to exon 5 (see Fig. S4). This involved co-injecting one-cell stage embryos with CRISPR-Cas9 RNP complexes containing two unique sgRNA sequences that flanked this region.

RNP complexes were generated using in vitro transcribed sgRNA oligonucleotides as previously described[74]. Briefly, double-stranded DNA oligonucleotides for sgRNA synthesis were made by PCR amplification using a constant oligonucleotide (AAAGCACCGACTCGGTGCCACTTTTTCAAGTTGATAACGGACTAGCCTTATTTTAACTTGCTATTTCTAGCTCTAAAAC), primers GCGTAATACGACTCACTATAG and AAAGCACCGACTCGGTGCCAC, and the variable target sequence noted below.

Upstream sequence before transcription start site:
GCGATTTAGGTGACACTATAGCTGTCTTTGGCACAAACAAGTTTTAGAGCTAGAAATAGC

Downstream sequence following exon 5:
GCGATTTAGGTGACACTATAGAGGAGCTCTGTTATCCGGGGTTTTAGAGCTAGAAATAGC

PCR amplification was performed by first combining 1 µL constant oligo (1 µM), 1 µL variable oligo (1 µM), 25 µL Quick-load 2x Master Mix (New England Biolabs, #M0271L), and 23 µL water. Following a first round of amplification (45 sec at 95℃, 30 sec at 60℃, 20 sec at 72℃), 2.5 µL of primer solution was added (10 µM each). Amplification was then performed by completing 34 cycles of amplification (15 sec at 95℃, 30 sec at 60℃, 20 sec at 72℃) and a final extension for 5 minutes at 72℃. DNA was gel purified using the QIAquick gel extraction kit (Qiagen, #28704) following protocol directions. The resulting double-stranded DNA contains four main parts: a three base-pair 'clamp' at the 5' end to stabilize the double-stranded sequence, a SP6 promoter, a variable targeting sequence, and a Cas9 binding scaffold ([clamp][promoter][guide sequence][Cas9 binding sequence]).

Upstream sequence before transcription start site:
[GCG][ATTTAGGTGACACTATA][GCTGTCTTTGGCACAAACAA][GTTTTAGAGCTAGAAATAGCAAGTTAAAATAAGGCTAGTCCGTTATCAACTTGAAAAAGTGGCACCGAGTCGGTGCTTT]

Downstream sequence following exon 5:
[GCG][ATTTAGGTGACACTATA][GAGGAGCTCTGTTATCCGGG][GTTTTAGAGCTAGAAATAGCAAGTTAAAATAAGGCTAGTCCGTTATCAACTTGAAAAAGTGGCACCGAGTCGGTGCTTT]

The sgRNA was then generated by In-vitro transcription using the HiScribe In Vitro Transcription Kit, following protocol directions. The sgRNA was then purified using a Qiagen RNEasy Plus mini kit (Qiagen, #73404) and diluted to 25 µM.

RNP complexes were prepared by combining 10 µL Cas9 (IDT, 62.5 µM Alt-R S.p. Cas9 Nuclease V3, #1081058), 14.8 µL Cas9 dilution buffer [cite Hoshijima] (20 mM HEPES, 350 mM KCl, 10% glycerol), and aliquoting into 2 µL RNP samples. Injection mixtures were then prepared by combining Cas9 and sgRNA in approximately equimolar proportions (1 µL diluted Cas9, 0.5 µL upstream sgRNA, 0.5 µL downstream sgRNA) and diluting with an additional 3 µL water. The RNP complexes were allowed to form for at least 10 minutes and approximately 1nL was injected per one-cell stage embryos.

Following CRISPR-Cas9-mediated editing in embryos, two F0 founder individuals (one male, one female) were identified by PCR-based genotyping. The knockout founders were intercrossed and mixed allele homozygous mutants were identified. These homozygous F1 fish or wildtype siblings were incrossed to generate the mutant and wildtype F2 embryos that were used directly in all subsequent experiments.

#### Genotyping of Galvanin Knockout in Zebrafish

Initial validation of gene editing was achieved by PCR amplification of the target region from embryo lysates. Lysates were prepared by combining embryos with 30 µL 50 mM NaOH and heated to 95℃ for 20 minutes and then cooling to 4℃. The pH was then reduced by adding 3.3 µL 1 M Tris-HCl (pH 8) and stored at -20℃ prior to use. Two PCR amplification were performed per sample using the following mixtures: 1) Positive excision: 6.25 µL Quick-Load Taq 2X Master Mix, 0.25 µL of 10 µM forward primer CCACAGGAGGAAAATGTTGTGGC, 0.25 µL of 10 µM reverse primer GTCAATCTACGTTCCTACTCTCACC, and 4.75 µL water, and 2) intact target region edit: 6.25 µL Quick-Load Taq 2X Master Mix, 0.25 µL of 10 µM forward primer CCACAGGAGGAAAATGTTGTGGC, 0.25 µL of 10 µM reverse primer GTCCAGCAGGCTGTCAATCC, and 4.75 µL water. PCR samples were then prepared by combining 11.5 µL of each mixture with 1 µL embryo lysate and PCR amplification performed (30 sec at 95℃, 40 cycles of 30 sec at 95℃, 30 sec at 53℃, 30 sec at 68℃, and then a final elongation for 5 minutes at 68℃). The first primer set spans the excised allele (indicating successful deletion), while the second primer set spans the intact target region (indicating presence of the wild-type allele).

To validate germline transmission and identify stable Galvanin knockout lines, genotyping was performed on adult fish through finclipping. Here, adult zebrafish were anesthetized in E3 embryo medium supplemented with 168 mg/L Tricaine (MS-222; Sigma-Aldrich, #E10521). The anesthetized fish were transferred to a clean surface, and a small biopsy (~2–3 mm) of the caudal fin was excised using a sterile scalpel blade. Biopsied fin tissue was transferred to individual PCR tubes and lysates prepared following the same approach noted for embryos above. PCR-based genotyping of fin clip lysates was performed using the same two-primer strategy as for embryonic validation.

#### Generation of Galvanin-eGFP expressing Cell Lines

Human HL-60 cell lines:

Cell lines expressing Galvanin-eGFP and the charge-modified proteins were integrated by lentivirus follow the same approach as the CRISPRi work above, using vectors custom cloned by Epoch Life Sciences (Texas). The Galvanin (TMEM154) gene coding sequence (NCBI mRNA reference sequence NM_152680.3), with eGFP linked to the intracellular c-terminus using a GGGGSGGGGSGGGGSGS amino acid linker sequence. This was driven from a SFFV promoter and included the native signal sequence. In our work looking at localization of Galvanin-eGFP and membrane protrusion/retraction activity (Fig. 3), the construct was expressed in our CRISPRi cell line (only expressing dCas9-KRAB, without any sgRNA) with or without an additional myosin-mApple label (*59*). A second version of the Galvanin-eGFP construct with an HA tag at the c-terminus was also designed, with the sequence GGGGSGGGGSG**YPYDVPDYA** appended after the eGFP sequence (bold indicates the HA tag). This was expressed in the Galvanin knockout cell line.

For the charge-engineered constructs, we removed the GGGGSGGGGSGGGGSGS + eGFP sequence but kept the GGGGSGGGGSG**YPYDVPDYA** HA tag sequence at the c-terminus. The entire extracellular domain between the signal sequence and transmembrane domain was replaced with the charged-GFP sequences (*33*) and a linker sequence (-42*e*: XTEN linker, identical to that used in the dCas9 sequence; +9*e*: GGGGSGGGGSGGGG) connecting to the native transmembrane sequence. The charged-GFP sequences were codon-optimized for expression in human cells.

Preparation of lentivirus and Integration into undifferentiated cells was performed identically to that described above for the CRISPRi cell lines using a spinoculation protocol. Enrichment of fluorescent-positive cells was achieved using fluorescence activated cell sorting (FACS, Sony Biotechnology Inc., SH800).

MDCK cell lines:

A fluorescent fusion construct of Galvanin expressing eGFP at the 3’ end was similarly generated for integration into MDCK cells. This was synthesized by GenScript and delivered on a pUC57-Kan plasmid backbone. The pUC57-Kan TMEM154-eGFP was then subcloned into a PiggyBAC transposon backbone. Ligation products were transformed into NEB Stable competent cells (NEB, catalog #C3040H) and positive colonies were confirmed by sequencing.

For transfection, 50,000 MDCK-II cells were seeded per well in a 24-well plate and transfected ~18 hours later using Lipofectamine LTX (Thermo Fisher Scientific, catalog #15338100).
Six days post-transfection, cells were subjected to FACS (Sony Biotechnology Inc., MA-900) to isolate cells with varying expressions of eGFP into 96-well plates to derive monoclonal lines. Clonal lines were screened for stable eGFP expression and membrane localization via flow cytometry (Invitrogen Attune NxT) and fluorescence microscopy (inverted Nikon Ti2 with NIS Elements software and a Nikon Qi2 camera).

### Genome-Wide and Secondary CRISPRi Assays

#### Development of Device for Pooled Screens

Our screen strategy involved exposing cells to an electric field, to drive their migration through the pores of a track-etch membrane. The aim was to isolate subpopulations where gene knockdown altered the ability of cells to sense the electric field and perform electrotaxis. Enrichment or depletion of specific sgRNAs would be expected following enhancement or disruption of electrotaxis, respectively. We have previously used track-etch membranes to perform screens of chemotaxis and undirected migration (i.e. chemokinesis) (*18*). In order to scale up an in vitro electrotaxis setup to allow exposure of millions of cells to an electric field we designed a device for large scale electrotaxis. Here, our initial focus was to ensure suitable exposure to an electric field, stable media conditions, and that the plastics and glue used in the device were non-toxic to cells. Below we detail these aspects in the context of the devices used in the screen work of the main text. In general, we followed conventions used in smaller scale electrotaxis devices, using a 'salt bridge' made of agarose to isolate the electrode chambers from the cell chamber. We note that the device used in the secondary screen was modified following some computational analysis using the finite element analysis software COMSOL Multiphysics (v. 6.1, COMSOL Inc., Sweden) to better understand the electrical properties.

The screen device was designed to be modular, allowing us to reuse key components such as the bulky center cell chamber, while making other components such as the track-etch membrane insert a one-time use item. We used Fusion 360 (Autodesk, San Francisco, CA) to design a device that could be printed using a 3D printer (UltiMaker S5 with Air Manager and Material Station). All design files are available upon request. Transparent PLA filament (Ultimaker, #M-X6C-Y6J2) was used for each printed plastic component.
In Fig. S1A we show a rendering of the assembled device used in the genome-wide screen, while in Fig. S1B we show a cross section. Each end contains a reservoir for Silver/silver chloride (Ag/AgCl) electrodes, immersed in a PBS solution. Ag/AgCl electrodes were generated by placing 20 gauge sheets of silver (Rio Grande, #101920) into a bleach solution for 30 minutes. During the screen, electrodes were connected to a direct current power supply (PowerPac 1000, Bio-Rad) via alligator clips, with a target total current of 400 mA. This resulted in joule heating of the cell culture media that was sufficient to heat it to 37℃, though we note that the total current was reduced during the experiment, if needed, to keep the temperature from going above 37℃.

Moving inward in the cross-section of Fig. S1B, the next segment is the agar salt bridge with 2% agarose in PBS. Between the salt bridges and central block is an auxiliary media reservoir that permits recirculation of culture media (with 5% hiFBS) utilizing a peristaltic pump. This reservoir is isolated from the central block with a track etch membrane, which aids in isolating the culture media in the central block from the agar bridges. Lastly, the central block where the cell migration experiments were performed contains the migration module that has the track-etch membrane. The track-etch membranes (3 µm diameter pores, shiny side oriented upward where cells are added; Sigma Millipore, #TSTP04700) were adhered to printed plastic inserts with silicone adhesive sealant (LOCTITE SI 5011 CL non-corrosive RTV, Henkel #51387). During assembly, the mating surface of the migration module and its seat in the central block was coated with silicone high vacuum grease (DuPont; Fisher Scientific #146355D) to create a water-tight seal with the central block. The assembly was held together with a conventional bar clamp using two aluminum plates to distribute the clamping force. We used a custom molded PDMS gasket between the central block and the auxiliary media reservoirs.

The expected electric potential across the surface of the track-etch membrane and electric field strengths were calculated using the finite element analysis software COMSOL (Fig. S1C). A model of the fluid-filled region was generated using Autodesk Fusion 360 and imported into COMSOL. Analysis was performed using their electric currents module. Key parameters included the temperature (293.15K) and media conductivity (1.27 S/m), which was experimentally measured for our cell culture media using a conductivity meter (Mettler SevenMulti, Mettler Toledo). In our analysis, the electric potentials and electric field strengths are based on a total current of 400 mA, taken to match the experimental conditions used.

The device used in the secondary screen is similarly rendered in Fig. S2A and a cross-section is shown in Fig. S2B. While maintaining most of the features noted above, the orientation of the anode electrode holder was moved to be directly above the migration module. In this more recent implementation, we also modified our migration module, which is shown in Fig. S2C. Since the HL-60 neutrophils are poorly adherent, it was possible for cells to get swept away from the module. In this version, a second track-etch membrane (Sigma Millipore, 1.2 µm diameter, #RTTP14250) was included to ensure cells stayed within the migration module. Cells were loaded into the module via a small hole with a pipette which was then sealed with silicone vacuum grease. As shown in our COMSOL analysis (Fig. S2D), this new device geometry produced a more uniform electrical environment at the surface of the track-etch membrane.

#### Overview of Cell Collection and Experimental Replicates

For each CRISPRi cell migration experiment, cells were collected from three populations for gDNA extraction: On the day of each experiment, 5-day differentiated HL-60 neutrophil cells were collected and $3\times10^7$ cells were set aside as a reference sample. The other two populations were the fraction of cells that migrated through the membrane and the fraction of cells that remained on top of the membrane.

Regarding experimental replicates, the genome-wide electrotaxis experiment involved 5 experimental replicates. For the smaller scale CRISPRi screen, the electrotaxis screen was performed 10 times, while the undirected migration screen was performed 8 times.

Cell migration screens identified gene candidates by comparing the number of cells that migrated through the track-etch membrane pores with respect to the reference sample, and those that did not with respect to the reference sample. This resulted in two separate measurements per migration experiment.

#### Removal of Cell Debris and Dead Cells Prior to Cell Migration Assays

Pooled CRISPRi libraries were differentiated in 15 cm dishes (55 ml cell culture per dish). Cellular debris and dead cells were removed from the differentiated HL-60 cell suspensions prior to use by density gradient centrifugation. Briefly, cells were first spun down (10 minutes at 300 g) and resuspended in 10 mL PolymorphPrep (Cosmo Bio USA #AXS1114683), placed in the bottom of a 50 mL conical tube. Using a transfer pipette, 15 mL of 3:1 PolymorphPrep : RPMI media + 10% hiFBS was gently layered on top by dispensing along the walls of the tube. This was followed by layering another 14 mL of RPMI media + 10% hiFBS. Cells were spun at 700 g for 30 min with reduced acceleration and braking to reduce mixing. Live differentiated HL-60 cells were collected between the RPMI media and the 3:1 PolymorphPrep. RPMI media layers were diluted with one volume of RPMI media + 10% hiFBS, and spun down once more for 10 minutes at 300 g. Finally, cells were resuspended in 10 ml RPMI media with 5% hiFBS and counted using a BD Accuri C6 flow cytometer (live cells identified by their forward-scatter and side-scatter, which show a single population separate from dead cells or debris)..

#### Cell Migration Screen Assay

Migration screens used track-etch membranes with 3 µm pore sizes as noted above in the device design. For each experiment, 20 million cells were added to the top of a track-etch membrane in the migration module. For undirected migration, devices were placed in a 37℃ incubator. For electrotaxis screens, experiments were performed on the lab bench, with joule heating from electrical stimulation maintaining the media at 37℃ as noted above. Following incubation for the required time (8 hours for undirected migration and two hours for electrotaxis experiments), the migration modules were removed from the central block. To ensure more complete recovery of the migratory cells, the bottom side of the track-etch membrane was gently scraped using a cell-scraper (Celltreat, #229310) to dislodge any cells remaining on the membrane surface. The migratory cells (bottom reservoir in central block) and retained cells (top of the migration module reservoir) were separately collected. Cells were spun down and washed with 1 mL PBS. Cells were spun down once more, with the PBS removed, and frozen at -80℃ for later genomic DNA extraction.

#### Quantification of sgRNA from CRISPRi libraries and Gene Identification

Genomic DNA (gDNA) was isolated using QIAamp DNA Blood Maxi ($3x10^7$ - $1x10^8$ cells) or Midi ($5x10^6$ - $3x10^7$ cells) kits following protocol directions (Qiagen, #51192 and #51183). gDNA precipitation was then used to concentrate the DNA. Briefly, salt concentration was adjusted to a 0.3 M concentration of ammonium acetate, pH 5.2 and 0.7 volumes of isopropanol were added. Samples were centrifuged for 15 minutes at 12,500 g, 4℃. Following a decant of the supernatant, the gDNA was washed with 10 mL 70% ethanol and spun at 12,500 g for 10 minutes, 4℃. The samples were washed in another 750 µL 70% ethanol, spun at 12,500 g for 10 minutes, 4℃, and decanted. The pellets were allowed to air-dry prior to resuspending them in water. The gDNA concentrations and purity were determined by UV spectroscopy.

The sgRNA sequences from each gDNA sample were PCR amplified for sequencing following protocols provided by the Broad Institute's Genetic Perturbation Platform. Briefly, gDNA samples were split across multiple PCR reactions, with 10 µg gDNA added per 100 µL reaction: 10 µL 10x Titanium Taq PCR buffer, 8 µL dNTP, 5 µL DMSO, 0.5 µL 100 µM P5 Illumina sequencing primer, 10 µL 5 µM P7 barcoded Illumina sequencing primer, and 1.5 µL Titanium Taq polymerase (Takara,# 639242). The following thermocycler conditions were used: 95℃ (5 minutes), 28 rounds of (95℃ (30 s) - 53℃ (30 s) - 72℃ (20 s)), and a final elongation at 72℃ for 10 minutes. PCR products (expected size of ~360 bp) were gel extracted using the QIAquick gel extraction kit (Qiagen, #28704) following protocol directions. After elution, samples were further cleaned up using isopropanol precipitation. Here, 50 µL PCR DNA samples were combined with 4 µL 5M NaCl, 1 µL GlycoBlue coprecipitant (ThermoFisher Scientific Technologies,# AM9515), and 55 µL isopropanol. Samples were incubated for 30 minutes and then centrifuged at 15,000 g for 30 minutes. The resulting pellet was washed twice with 70% ice-cold ethanol and resuspended in 25 µL of Tris-EDTA buffer. Illumina 150bp paired-end sequencing was performed by Novogene Corp. (Sacramento, CA).

Sequence reads were quality filtered by removal of reads with poor sequencing quality and reads were associated back to their initial samples based on an 8 bp barcode sequence included in the P7 PCR primer. The 20 bp sgRNA sequences were identified and mapped to gene targets using a reference file for the genome-wide CRISPRi library (*55*). As noted in the overview section above, two $\log_2$ fold-change values were calculated from the sequencing counts from each pooled screen experiment: each enriched sample (top or bottom reservoir collected cells) compared against the reference sample collected prior to the experiment. Since we expect these two measurements to be inversely correlated, the $\log_2$ fold-change values from the less-migratory population collected in the top reservoir were multiplied by -1. This allowed the two sets of $\log_2$ fold-change values to be compared directly, and we averaged across all such measurements. Reported $\log_2$ fold-changes represent averages across median-normalized replicate measurements from the multiple experiments performed. Here, a pseudocount of 32 was added to the sgRNA counts to minimize erroneously large fold-change values in cases of low library representation (*60*). $\text{Log}_2$ fold-changes were also scaled to have unit variance prior to averaging across individual experiments. P-values were determined by performing permutation tests (*61*) between the calculated $\log_2$ fold-change values for each gene target and our set of control sgRNA $\log_2$ fold-change values. Adjusted p-values for multiple comparisons were determined using the Benjamini–Hochberg procedure (*62*).

#### Image Acquisition

All microscopy-based image acquisition was performed using microscope setups operated by MicroManager (v. 2.0) (*63*). Details of the microscope configurations are provided below for each cell migration assay.

### Three-Dimensional Migration in Collagen Extracellular Matrices

#### Experimental

HL-60 neutrophils and EL4 T cells were prepared for microscopy as previously described (*18*, *54*). For HL-60 neutrophils, we additionally removed dead cells and debris following the Polymorphprep protocol described above. Briefly, following clean up, $3x10^5$ cells were collected, resuspended in 1 mL L-15 + 10% hiFBS media containing 1 µg/mL DNA stain Hoechst 33324 and incubated at 37℃ for approximately 15 minutes. During incubation with Hoechst stain, a 200 µL collagen aliquot was prepared: 6.5 µL 10x PBS, 12.5 µL 0.1 M NaOH, 111 µL L-15, and 20 µL hiFBS were combined on ice, with 50 µl 3 mg/mL collagen added and mixed just prior to resuspension with cells. The cell suspension was spun down and resuspended in the mixture, with a target concentration of 0.75 mg/mL collagen. This was immediately added to the channel of an Ibidi µ-Slide I (Ibidi, #80106). After 1 minute incubation at room temperature with the slide inverted, the channel slide was placed coverslip side down and incubated at 37℃ for gel formation for 17-18 minutes. Inverting the slide helps counter cell sedimentation as the gel sets. The media reservoirs were then filled with 0.75 ml L-15 media containing 10% hiFBS. Imaging was performed within approximately 30 minutes after the gel set.

Cells were exposed to an electric field using a custom device similar to that previously described (*23*, *64*). We designed a platform that would hold the Ibidi sample slide and electrode reservoirs filled with a PBS solution and fit into a 96-well plate microscope stage insert. Salt bridges were 3D printed and filled with 2% agarose in PBS to connect the electrode reservoirs to the Ibidi media reservoirs. All device pieces were printed using transparent PLA filament (Ultimaker, #M-X6C-Y6J2), 3D files available upon request. Ag/AgCl electrodes were produced by immersing strips of silver foil (0.127 mm thick; Alfa Aesar, Cat#11440-GW or 30-Ga .999 fine silver from Rio Grande, #101930) in bleach for 30 minutes and then rinsing several times with water and 70% ethanol. The electrodes were connected to a direct current power supply (Keithley Instruments #2200-72-1 or B&K Precision #BK9184B-ND) via alligator clips. The power supply was used in constant current mode to maintain a constant current density and voltage was measured using a multimeter by immersing platinum electrodes at each side of the Ibidi channel.

Cells were imaged at 37℃ on a Nikon Ti2 inverted microscope, equipped with a piezo-z stage (Applied Scientific Instruments PZ-2300-XY-FT), a Yokogawa CSU-W1 spinning-disk confocal, and iXon EMCCD camera (Andor). Imaging was performed with a 20x 0.95 NA water objective lens using sequential brightfield and epifluorescence illumination (405 nm laser; CSU-W1 Penta Dichroic; Emission: Chroma ET450/40m). For each sample, a 30 min time-lapse movie was acquired with 60 s intervals. A z-stack was acquired over 300 µm with acquisitions every 3 µm. In general experiments were performed over three different days using freshly prepared slides.

#### Cell Tracking and Quantification

Cell tracks were extracted from the DNA channel of time-lapse microscopy images using the TrackMate package (v. 7.12.2) (*65*) in FIJI (v. 1.54) (*66*). Any stage drift was corrected using a small number of non-motile cells present in the collagen gel. These were taken as fiducial markers, with x, y, z drift correction performed to maintain their non-moving position. Cell track information, including position and time, was aggregated into a table using pandas Python package (v. 1.4.4) (*67*). For the HL-60 Galvanin knockout expressing Galvanin-GFP (i.e. the genetic rescue line), we note that cells were not sorted prior to migration experiments and there was a small cell subset (approximately 20%) that were not Galvanin-GFP positive. Prior to each acquisition, a single z-stack was taken of the GFP channel (488 nm) and this data was used to filter out non-fluorescent cells. For EL4 cells, due to poor signal in the fluorescence channel, tracking by TrackMate was performed on z-max-projected images, obtaining only the x-y trajectory information.

Cell migration speeds were calculated by subsampling the three-dimensional track vectors every 180 seconds. Calculations were performed on an individual cell basis, with an average migration speed calculated across all time intervals in the 30 minute video. The average speed along the average speed along the electric field vector was similarly calculated, but in this case we consider the x-component (i.e. electric field direction) of the track vectors. Comparisons of the Galvanin knockout and Galvanin-GFP rescue with the wild-type cells were performed using the two-sided Mann-Whitney U nonparametric test scipy.stats.mannwhitneyu() in the SciPy Python package (v. 1.9.1) (*68*). Cell tracks longer than 16 minutes were included in this analysis.

The compass autocorrelation involved first calculating the cosine angle ($\cos\theta$) between the migration track vector and the electric field vector at each 180 second interval (see Fig. S3B for a schematic illustration). For each cell, this resulted in an array $D(t)$ of $\cos\theta$ values over the 30 minute video. A normalized non-overlapping autocorrelation was calculated by iterating over each possible time lag. Here we compute the sum of products between non-overlapping pairs of $\cos\theta$ values,

$$S(\tau) = \sum_{i=0}^{N-\tau-1} D(i\cdot(\tau + 1)) \cdot D(i\cdot(\tau + 1) + \tau).$$

We then normalize by the square root of the product of the sum of squares of the corresponding non-overlapping segments of the signal at the current and lagged positions,

$$Compass\ Autocorrelation\ (\tau) = \frac{S(\tau)}{\sqrt{\sum_{i=0}^{N-\tau-1} D(i\cdot(\tau+1))^2 \cdot \sum_{i=0}^{N-\tau-1} D(i\cdot(\tau+1)+\tau)^2}}.$$

Only long cell tracks (27 minutes or longer) were used in the autocorrelation analysis.

### Two-Dimensional Migration

#### Zebrafish Keratocytes: Cell Isolation, Slide Preparation, and imaging

Keratocyte cells were isolated from whole 2 days post fertilization embryos as previously described [75,76]. Briefly, for each preparation 10 embryos were dechorionated and anesthetized with 160 mg/ml Tricaine. Larvae were then washed twice in PBS and incubated in 400 μL cell dissociation buffer (Gibco #13151014) at 4°C for 30 min on a tube rotator. Cell dissociation buffer was then removed and replaced with 200 μL 0.25% trypsin and 1 mM EDTA for 15 minutes at 28℃. The trypsin was quenched using 200 μL fetal bovine serum supplemented with antibiotic-antimycotic and pipetted to help dissociate cells further. Cell nuclei were stained by incubating cells with 1 μg/mL DNA stain Hoechst 33324 for 15 minutes, concentrated by centrifugation for 5 minutes at 500 g, and resuspended in 100 μL imaging media (L-15 media supplemented with 10% fetal bovine serum and antibiotic-antimycotic). The cell suspension was the added to an Ibidi μ-Slide I (Ibidi, #80106) and incubated at 28℃ to allow cells to adhere. Finally, debris and non-adhered cells were removed by adding fresh imaging media in one well of the slide and aspirating the media from the opposite well. This was repeated three times and then another 800 μL imaging media was added to each reservoir well of the slide. Cells were imaged at 26℃ on two inverted microscopes (Nikon Ti Eclipse), with epifluorescence illumination using a similar configuration as noted below for HL-60 neutrophils. Imaging was performed with an 20x air objective lens (Nikon 20x 0.75 NA plan apo phase contrast) using sequential brightfield and epifluorescence illumination and captured on an iXon EMCCD camera (Andor) and 1 minute imaging intervals.

#### MDCK epithelial cells: Slide Preparation and Imaging

MDCK cells were seeded in Ibidi μ-Slide I at low density to monitor the electrotaxis behavior of single cells and small cell clusters. Since cells adhere quite strongly to each other, trypsinized cells were vortexed and pipetted multiple times to ensure a single cell suspension before seeding. For each slide preparation, 80,000 cells were resuspended in 1 mL of DMEM media (Gibco, #11885084) supplemented with 10% FBS, 100 U/mL penicillin, and 100 ug/mL streptomycin (Gibco, #15140122) and added to the slide. Cells were incubated at 37°C for approximately 3 hours to allow cells to adhere to the coverslip. Prior to imaging, the media was replaced with fresh media containing 1 μg/mL DNA stain Hoechst 33324 and incubated for 15 minutes at 37°C. Following two washes to remove the residual DNA stain, 0.8 mL of media was added to both reservoir wells of the slide. Sample preparations were imaged at 37℃ using two inverted microscopes (Nikon Ti Eclipse), with epifluorescence illumination using a similar configuration as noted below for HL-60 neutrophils. Imaging was performed with an 20x air objective lens (Nikon 20x 0.75 NA plan apo phase contrast) using sequential brightfield and epifluorescence illumination and captured on an iXon EMCCD camera (Andor) and 5 minute imaging intervals. Imaging was performed over 4 days, with two sample replicates per condition and cell line. For one replicate, the total imaging time was two hours, while for the second replicate it was extended to three hours. We note that in several samples drift was observed in the first several frames. These were excluded from the data analysis.

#### HL-60 Neutrophils: Slide Preparation and Imaging

Cells were resuspended to a concentration of 2x10$^{6}$ cells/ml in L-15 Medium with 10% hiFBS. With an ibiTreat µ-Slide-I (Ibidi, #80106) placed on a 37℃ heat block, 100 µL of cells were added to the channel and incubated for 5 minutes to allow them to sediment and loosely adhere to the bottom coverslip. Warm, liquid 2.5% UltraPure low melting point agarose (Invitrogen, #16520-050) in L-15 with 10% hiFBS was added to one channel reservoir and then centrifuged at 28-37℃ for 5 min at 700g. This was performed in a custom 3D-printed swinging bucket rotor plate adaptor (design available upon request). Immediately after centrifugation, the slide was incubated on a 20℃ cooling plate (STIR-KOOL, Ladd Research, #SK-12D-AW) for 5 minutes to gel. Excess solidified agarose was observed in the media reservoirs of the Ibidi slide and removed with a pipet tip. L-15 media with 10% hiFBS was added to both reservoirs.

For under agarose experiments with Latrunculin A, cells were prepared as for other under agarose experiments, except prior to addition of cells to the µ-Slide I channel, 254 µm diameter 8 lb Maxima Ultragreen monofilament fishing line (West Marine, Seattle) was threaded into the channel so that it stuck out on either end. After the agarose solidified, 25 µM Latrunculin A (Invitrogen, # L12370) in complete media was added to each reservoir. The monofilament line was removed, leaving a tunnel free of agarose the length of the Ibidi channel. Excess agarose was removed from the reservoirs. This thin tunnel, free of agarose, allowed flow of the media containing Latrunculin A along the channel length. This is important since it shortens the distance the drug must diffuse through the agarose.

Cells were imaged at 37℃ on an inverted microscope (Nikon Ti Eclipse), with epifluorescence illumination using a Lambda 721 (721CUBE-480, Sutter Instruments) and standard GFP filter cube (Chroma, ET-EGFP #49002; Ex: ET470/40x; Dichroic: T495LPXR; Em: ET525/50m). Imaging was performed with an oil 60x objective lens (Nikon 60x 1.4 NA plan apo phase contrast) using sequential brightfield and epifluorescence illumination and captured on an iXon EMCCD camera (Andor). For non-drug treated samples, imaging was performed at 5 second intervals, while for the Latrunculin A, imaging was performed at 10 second intervals. Exposure of cells to an electric field was performed as described in section ‘Three-Dimensional Migration in Collagen Extracellular Matrices,’ above. In the case of Supplemental Movie 1, we used a custom design where instead of the Ibidi slide, cells were added to a glass coverslip and overlaid with a 510 µm thick 1% agarose/L-15/10% hiFBS gel as previously described (*54*). For this movie, imaging was performed using a 100x objective lens (Nikon 100x 1.45 NA plan apo phase contrast with 1.5x additional magnification applied).

#### HL-60 Neutrophils: Quantification of Peripheral Galvanin-GFP Signal and Membrane Velocity

Segmentation of cells was performed using the phase images. Here, an image that is just the agarose background was subtracted pixelwise from the phase image, then the contrast was adjusted by renormalization. A Sobel discrete differentiation operator was then applied to the image, allowing us to identify the cell edge using a threshold cutoff, which was used to generate a binary mask of each cell. The fluorescence images were flat field corrected. The resultant segmentation masks and fluorescence images were rotated 90 degrees to the left and input to the ADAPT Plugin in FIJI (v.1.111) (*27*), with smoothing sigma of 2, a 2 μm cortex, and with 8 erosions. The signal kymograph and the velocity kymograph were generated by averaging across kymographs from individual cells. Only cells that were in frame and had no collisions nor segmentation errors over the collection period were included in the analysis.

We quantified the cross–correlation,

$$C(\tau) = \frac{1}{m}\sum_{t=0}^{m-1}\frac{(GFP(t) - \overline{GFP}) \cdot (vel(t+\tau) - \overline{vel})}{\sigma_{GFP} \cdot \sigma_{vel}},$$

which computes the correlation between the average Galvanin-GFP signal ($GFP(t)$) and membrane velocity ($vel(t)$) at each time point ($t$). Analysis was performed using data from the first 10 minutes, as a function of different time lags ($\tau$). In the equation above, $\overline{GFP}$ is the average of $GFP$(t) and $\overline{vel}$ is the average of $vel(t)$ over the ten minute interval, while $\sigma_{GFP}$ and $\sigma_{vel}$ are their standard deviations, respectively. $m$ is the length of the data arrays. This analysis was performed at both the anodal side (i.e. what becomes the cell rear; averaging across positions +120 to +240 of the kymographs) and cathodal side (i.e., what becomes the cell front; averaging across positions -60 to +60 of the kymographs) to look at the onset of the directional response.

To better assess statistical significance, we also calculated a confidence interval for the maximum cross-correlation value of randomly permuted data based on the input data. Using a bootstrapping approach (*48*), we generated 2000 bootstrap samples by randomly permuting the values in each of the $GFP(t)$ and $vel(t)$ arrays independently. For each bootstrap sample, we calculated the maximum cross-correlation value and a 99% confidence interval was estimated by determining the 1th and 99th percentiles of the distribution of maximum cross-correlation values from the bootstrap samples.

#### HL-60 Neutrophils: Quantification of $v_E$/D Ratio and Estimation of Charge

The distribution of Galvanin in an electric field can be described by a combination of diffusion and drift processes. Under our experimental conditions, we are specifically interested in describing the expected profile of Galvanin on the approximately flat two-dimensional bottom surface near the coverslip. Under our imaging conditions (≈ 800 nm axial resolution), we expect the measured fluorescence intensities to reflect fluorescence predominantly at this surface and we treat our biological system as a flat two-dimensional surface. Further, in our analysis we only consider the Galvanin profile after long exposure to an electric field (≳ 3 minutes), where the biased Galvanin distribution appears stable. Under this steady-state condition, the effects of drift and diffusion balance. With the electric field oriented along a single axis (taken to be the $x$ axis in the description below), under steady state we expected the protein concentration to decay exponentially along this axis (*31*). The probability distribution can be described by

$$P(x,y) = A\,e^{\frac{v_E}{D}x},$$

where $v_E$ is the electrophoretic drift velocity and D is the diffusion coefficient. Here, $A$ is a normalization constant that is determined by boundary conditions and the amount of Galvanin in the plasma membrane.

The electrophoretic drift velocity $v_E$ is dependent on the force $F_E$ acting on the charged protein and drag on the particle,

$$v_E = \frac{F_E}{\zeta},$$

where $\zeta$ is the drag coefficient. The force $F_E$ is determined by the Coulombic interaction between the protein with charge *q* (or *z* elemental charges $e^-$) and the electric field, given by,

$$F_E = qE\,f(\kappa) = ze^-E\,f(\kappa).$$

The factor $f(\kappa)$ accounts for counterion screening that will result in a reduced effective (i.e. measurable) charge that depends on the Debye screening length $\kappa^{-1}$. From the Einstein relationship, we can also relate the drag coefficient $\zeta$ to Galvanin's diffusion coefficient through

$$\zeta = \frac{K_B T}{D},$$

where $K_B$ is the Boltzmann constant, and T is the temperature. This allowed us to write,

$$\frac{v_E}{D} = \frac{ze^-E\,f(\kappa)}{K_B T},$$

which provides us with an expected relationship between the drift velocity, diffusion coefficient, protein charge, and the strength of the electric field (*23*).

We approximate the counterion screening using the Debye–Hückel approximation that describes the electric double layer of counter ion screening. In general, we expect a tightly bound layer of counter ions to move with the charged protein, whose size is larger than the screening length scale and which will result in an effective charge that is less than the bare charge (*69*). In order to estimate the screening factor $f(\kappa)$, we use the approximation that the electric potential $\phi(r)$ at a distance $r$ from a charge will decay exponentially (*70*),

$$\phi(r) = \phi \cdot e^{-\kappa r}.$$

The effective charge will be directly proportional to the electric potential,

$$\phi(r) \propto q(r) \propto q\,e^{-\kappa r},$$

because of the charge shielding. Taking the screening length to be the Debye length, $r = \kappa^{-1}$, we expect an approximate effective charge given by $0.37ze^-$. The estimated number of elemental charges reported in the main text figures is given by $z$.

As shown in Fig. 4A, we quantified the $v_E/D$ ratio across individual cells. Here we selected a region in the middle of each cell approximately 10-20 pixels thick, adjusted based on the cell shape or presence of bright fluorescence features. Background cellular autofluorescence was first subtracted from the intensity values, determined by measuring the intensity near the cathodal edge of cells exposed to a 500 mV/mm electric field and where no membrane-localized Galvanin-GFP signal was observed. The corrected fluorescence intensities were then log-transformed and the the $v_E/D$ ratio was determined from a linear fit of the data using the polyfit() function in the NumPy Python package (v. 1.21.6).

#### HL-60 Neutrophils: Quantification of Galvanin's Diffusion Coefficient

Galvanin quickly returns to a uniform distribution through diffusion in the plasma membrane after removal of the electric field. The fluorescence signal of Galvanin-GFP at the cell periphery (Fig. S4A) represents a one-dimensional slice of this two-dimensional process. This fluorescence is peaked at the anodal side of the cell and can be well described by a Gaussian distribution (Fig. S4B). We take the method used by the PIPE (photo-converted intensity profile expansion) approach (*71*), which is to estimate the diffusion coefficient by quantifying the spatial expansion of our fluorescence signal over time. This involves fitting the fluorescence profile to a Gaussian distribution, with the expectation that the squared-width of the Gaussian distribution will increase linearly over time. We expect this to only be valid at early time frames since the width of our fluorescence peak is already quite wide relative to the length scale of the entire cell and therefore only consider the 1-1.5 minutes following removal of the electric field (Fig. S4C).

To quantify the fluorescence around the periphery we used custom code with the Shapely Python package (v. 2.0.4) to convert our cell masks into a cell line edge, which was used to generate 300 equally spaced points along the cell boundary. For each point, we then generated a line segment orthogonal to the cell edge and calculated the maximum fluorescence intensity along that line segment to generate a fluorescence profile around the cell periphery. We fit this intensity profile to a Gaussian function using the curve_fit() function in the SciPy Python package (v. 1.9.1). Following the approach of ref. (*71*), the intensity data $I(k, t)$ was fit to a Gaussian function of form,

$$I(k, t) = A \cdot e^{\frac{-(k-\bar{k})^2}{S^2(t)}},$$

where A is a normalization factor related to the fluorescence intensities, $k$ is the distance along the periphery of the cell, $t$ is the time, $\bar{k}$ is the mean value where the intensity is peaked, while the variance $\sigma^2 = S^2(t)/2$. The diffusion equation follows an identical functional form, and we can relate the width of the profile to the diffusion coefficient, where $S^2(t) = S_0^2 + 4Dt$. $S_0$ is the initial width of the peak profile. The square widths $S^2(t)$ from these fits as a function of time were then fitted using the polyfit() function in the NumPy Python package (v. 1.21.6) to extract the slope and diffusion coefficient (slope = $4D$).

### Immunolabeling for Western Blots

Whole-cell protein lysates were collected from differentiated HL-60 cells for Western blot analysis. For each sample, $5x10^6$ cells were collected, washed once in ice-cold PBS, and resuspended in two volumes of 5x Laemmli SDS-PAGE sample buffer by weight (e.g. 40 µL for 20 mg cell pellet). Samples for heated to 98℃ for 5 minutes and then vortexed briefly prior to sonication with a bath type sonicator (Diagenode, #B01020001). Sonication was performed on their high power setting at 4℃ with five cycles of 30 seconds on and 30 seconds off. Samples were stored at -20℃ and re-heated to 98℃ prior to gel electrophoresis.

Samples were run on 12% polyacrylamide gels with a protein ladder (Bio-rad, #1610317) and and transferred to nitrocellulose membranes (Bio-rad, #1620233) by semi-dry transfer in buffer 10mM CAPS pH 11, 10% methanol. Transferred protein was confirmed and normalized using a reversible total protein stain kit (Pierce, #24580). Blots were then blocked in Tris-buffered saline with 0.1% Tween 20 (TBST) with 5% non-fat milk (spun at 2,500 g to remove milk precipitates). Blocking was performed for 30 minutes at room temperature. Protein loading and transfer efficiency was also assessed by staining the residual protein in the gel using Coomassie stain (0.006% Coomassie R250 with 10% acetic acid). Primary antibodies were diluted in TBST with 0.5% non-fat milk and incubated overnight at 4℃. Blots were washed with TBST for 30 minutes, with buffer exchanged every five minutes, and then stained with an HRP conjugated secondary antibody diluted in TBST with 0.5% non-fat milk. Following incubation for 60 minutes at room temperature, the blots were washed for 60 minutes in TBST, with buffer exchanged every five minutes. The blots were imaged with a digital gel documentation system (Azure c600), allowing for detection of the secondary HRP antibody detected using a chemiluminescence peroxidase substrate kit (Sigma, #CPS1A120).

#### Antibodies and Dilution Information

Primary antibody: Polyclonal anti-humanTMEM154 (1:500; Proteintech, #24812-1-AP). We note that the polyclonal antibody was found to produce substantial background when blotted against our differentiated HL-60 neutrophil samples. A secondary-only control blot confirmed the background resulted from the primary antibody. To reduce background, prior to probing blots, the diluted primary antibody sample was preabsorbed with total protein from our differentiated HL-60 Galvanin knockout sample bound to nitrocellulose. Briefly, 10 µL of SDS-PAGE sample was diluted with 40 µL water and 2 µL 3 M KCl added to precipitate potassium SDS. The precipitant was removed by centrifugation and the supernatant was diluted in 3 mL TBST and incubated with a piece of nitrocellulose of approximately 50 $cm^2$ at room temperature for one hour. Following several washes in TBST, the nitrocellulose was blocked using TBST with 5% non-fat milk for one hour at room temperature. The diluted polyclonal anti-TMEM154 antibody (3 mL total, 1:500 diluted in TBST with 0.5% non-fat milk) was then incubated with this nitrocellulose for one hour for preabsorption prior to use for Western blot.

Secondary antibody: HRP-linked anti-rabbit IgG (1:3000; Cell Signaling, #7074S).

### Supplemental Figures and Figure Captions

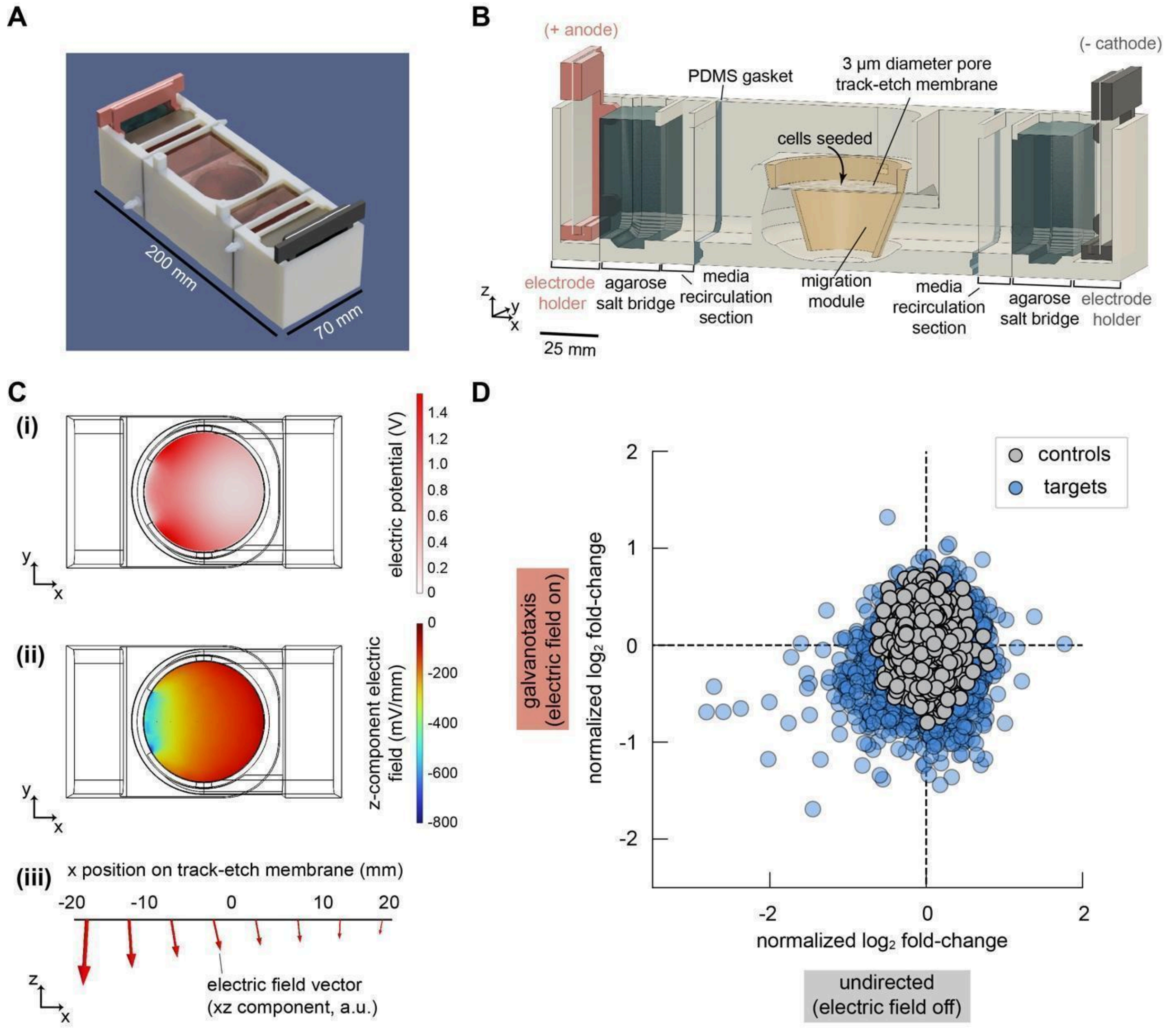


#### Figure Supplement 1

**Cell separation device for genome-wide CRISPRi screen. A.** Rendered image of assembled electrotaxis/ galvanotaxis screen device. **B.** Cross-section of screen device. Key components include Ag/AgCl electrodes and agarose salt bridges that isolate electrodes (in PBS) from region contain cells (in RPMI culture media with 5% hiFBS), and the center migration module containing a track-etch membrane (3 µm diameter pore size). Media was recirculated using the luer fittings connected to the 'media recirculation section', which recycled into a common beaker containing an additional liter of culture media. Colors are added to better distinguish different parts and features. **C.** Finite element analysis using COMSOL Multiphysics software was used to determine the electrical environment at the surface of the track-etch membrane. (i) Electric potential (voltage) across the surface of the membrane plane, taken relative to the smallest value calculated. (ii) Z-component of the electric field vector. (iii) Cross section at the plane of the track-etch membrane, showing the z-x component of the electric field vector (with the y-position centered on the membrane). **D.** Summary of genome-wide library screen targeting 18,901 genes for knockdown. Data points show the normalized $\log_2$ fold-change averaged across three sgRNAs per gene across independent experiments (5 screen replicates for electrotaxis and 4 screen replicates for undirected migration). The undirected migration screen data comes from prior work (*18*). Control values were generated by randomly selecting groups of three control sgRNAs.

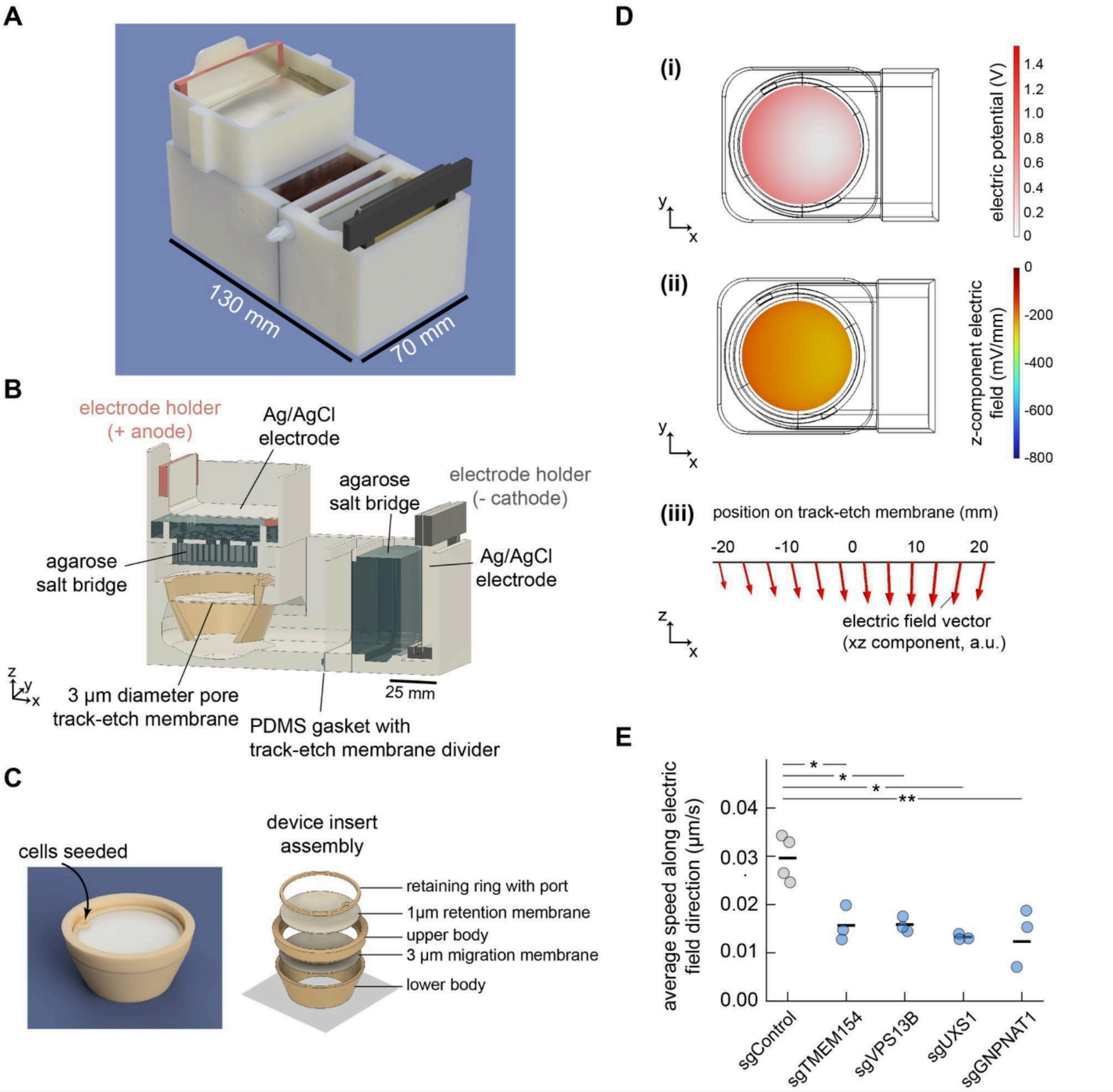


#### Figure Supplement 2

**Cell separation device for secondary, focused CRISPRi screen. A.** Rendered image of improved electrotaxis/ galvanotaxis screen device. **B.** Cross-section of screen device. Key components include Ag/AgCl electrodes and agarose salt bridges that isolate electrodes (in PBS) from region contain cells (in RPMI culture media with 5% hiFBS), and the migration module insert containing a track-etch membrane (3 µm diameter pore size). Peristaltic pumps were used to recirculate media into a beaker containing an additional liter of culture media, via the fittings shown, to help maintain uniform buffer conditions. Colors are added to better distinguish different parts and features. **C.** Migration module insert. Left: rendered drawing of assembled insert. Right: The different pieces are shown in an expanded view. Each piece was printed separately and assembled with the track-etch membranes. The upper track-etch membrane with the retaining ring and cell seeding port was included to ensure the cells remained in place during handling of the device. **D.** Finite element analysis using COMSOL Multiphysics software was used to determine the electrical environment at the surface of the track-etch membrane. This was done to improve the uniformity of the electrical environment relative to the device used in our initial screen. Plots show the (i) Electric potential (voltage) across the surface of the membrane plane, taken relative to the smallest value calculated, (ii) Z-component of the electric field vector, and (iii) Cross section at the plane of the track-etch membrane, showing the z-x component of the electric field vector (with the y-position centered on the membrane). **E.** Quantification of average speed along the electric field direction during migration in a collagen gel with exposure to an electric field (300 mV/mm). Individual data points represent the average across cells from a single collagen preparation and the average is indicated by the horizontal line representing a total of 500-850 cells per cell line). A Tukey's range test was performed to make multiple comparisons across the cell lines, with only the sgRNA knockdown versus sgRNA controls found significant (* $p$-value $< 0.05$, ** $p$-value $< 0.01$).

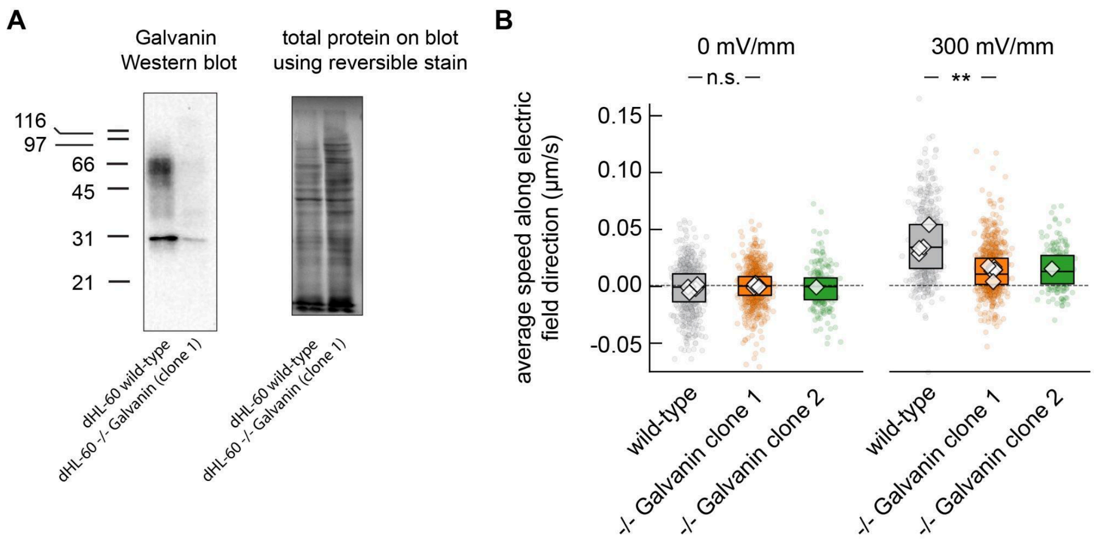


#### Figure Supplement 3

**Validation of Galvanin knockout in HL-60 cells. A.** Western blot confirms loss of Galvanin protein expression. Left: blot using a polyclonal antibody against Galvanin. We find a near-complete loss of signal in the Galvanin knockout (clone 1) when compared to wild-type HL-60 neutrophils. The smear is expected from variable protein glycosylation. Right: total protein stain on the blot using a reversible protein staining kit. Images are representative of Western blots performed in triplicate. **B.** Average speed parallel to the electric field direction calculated based on nuclear tracking with 3 minute time intervals. Individual colored scatter points represent average values of individual cells, while the boxplot extends from the first to third quartiles with a line at the median cells. The white diamonds represent average values across cells from experimental replicates). A significant difference is noted between the Galvanin knockout clone 1 and wild-type cells when exposed to a 300 mV/mm field (p-value=0.009, two-sided Mann-Whitney U test). A similar test was not performed for clone 2 since only a single experimental replicate was performed. However, it is noted that when comparing individual cell speeds, a significant difference is found between Galvanin knockout clone 2 and wild-type cells (p-value<0.0001, two-sided Mann-Whitney U test). Clone 1 is used throughout the manuscript, while clone 2 was also genotyped to confirm gene disruption. Each experimental condition represents average speed values from 200-900 cells.

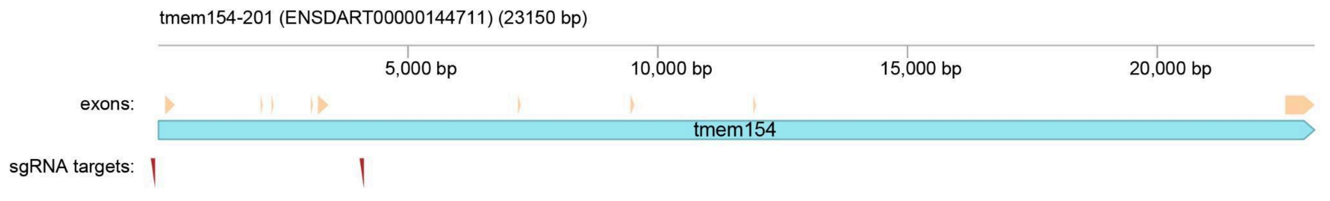


**Figure Supplement 4**

Galvanin knockout strategy in zebrafish. The *Danio rerio* (zebrafish) coding sequence for Galvanin (TMEM154) is shown in blue, with exons identified in yellow. The sgRNA target sites are identified in red (GCTGTCTTTGGCACAAACAA for sequence upstream of exon 1, and GAGGAGCTCTGTTATCCGGG for the sequence downstream of exon 5).

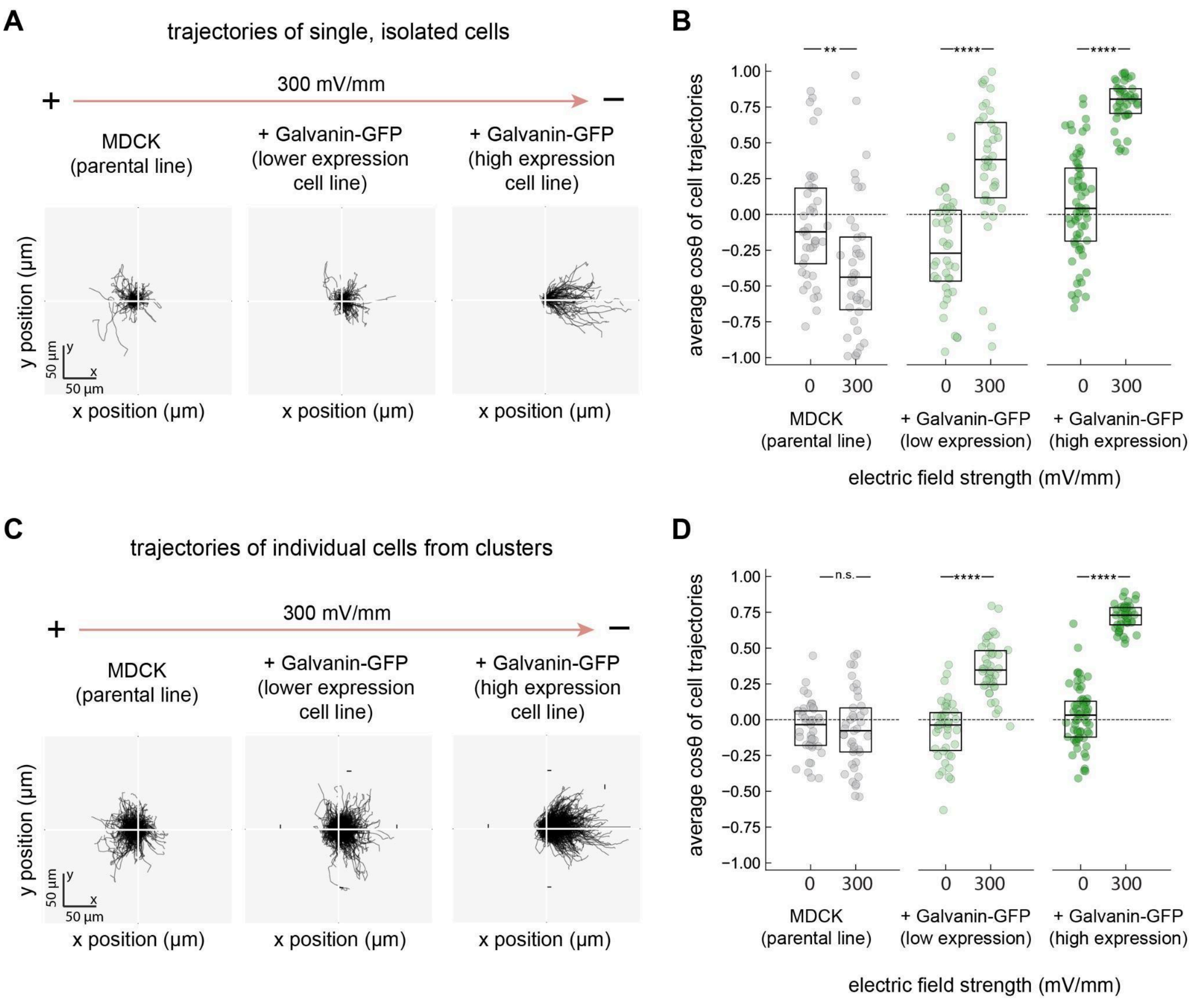


#### Figure Supplement 5

**Galvanin expression in MDCK cells is sufficient for cathodal electrotaxis of single cell and small cell clusters. A.** Relative cell trajectories of isolated, single MDCK cells exposed to an electric field (300 mV/mm) over three hours (5-minute imaging interval). A subset of trajectories is shown (n=100 per condition). **B.** Average cosine $\theta$ for isolated, single MDCK cells. Each data point represents the average across cells from a different imaging field of view (40 fields of view across two sample preparations, 157-404 cells per condition). **C.** Relative cell trajectories of cells in clusters exposed to an electric field (300 mV/mm) over three hours (5-minute imaging interval). Clusters were defined to have two or more cells in contact during imaging. A subset of trajectories are shown (*n*=100 per condition). **D.** Average cosine $\theta$ for small MDCK clusters. Each data point represents the average across cells from a different imaging field of view (40 fields of view across two sample preparations; 157-404 cells per condition). Data from panels **A**-**B** and **C**-**D** represent subsets of the data shown in Figure 4 of the main text. Asterisks indicate statistical significance between the 300 mV/mm and 0 mV/m conditions (** p-value=0.003; **** p-value < 0.0001, two-sided Mann–Whitney U test).

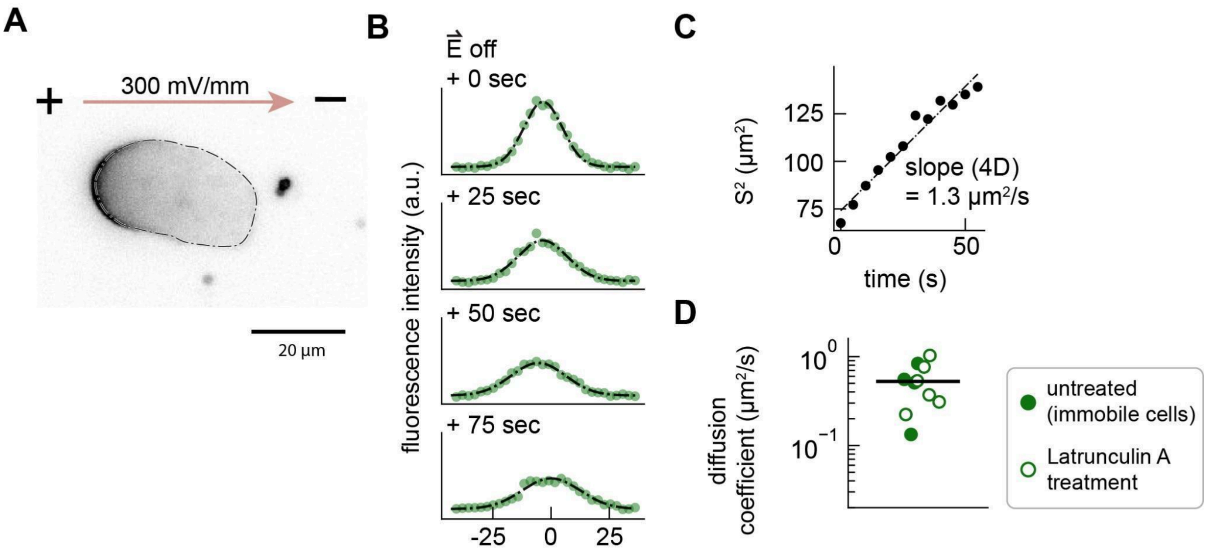


#### Figure Supplement 6

**Galvanin exhibits rapid membrane diffusion consistent with a single-pass transmembrane protein. A.** Example fluorescence micrograph of a cell exposed to an electric field (300 mV/mm) for 5 minutes. The dot-dashed line indicates the cell periphery used in the quantification of Galvanin-GFP fluorescence intensity. **B.** Plots show the decay in fluorescence intensity at the anodal side of the cell once the electric field is turned off, corresponding to a re-equilibration of Galvanin throughout the plasma membrane. Dot-dash line indicates a Gaussian fit of the fluorescence data. **C.** The squared width $S^2$ was used as a measure of the increase in the mean-squared displacement of the fluorescence signal over time, with an expected linear relationship between $S^2$ and 4D, where D is the diffusion coefficient of Galvanin-GFP. **D.** Estimation of diffusion coefficient. Data points represent measurements from individual cells following the analysis steps of A-C. The black line indicates the average across all measurements with an average value of 0.53 $\mu m^2/s$ (± 0.1 SEM).

### Captions for Supplemental Items

##### Supplemental Movie 1

**Galvanin-GFP biases toward the anodal side of the cell following exposure to electric field.** Epi-fluorescence microscopy movies of differentiated HL-60 neutrophils expressing Galvanin-GFP and exposed to an electric field of approximately 1000 mV/mm. Cells were confined by a 1% agarose gel (in L-15 media with 10% hiFBS) and imaged using an oil 100x 1.45 NA plan apo phase contrast objective with 1.5x additional magnification applied. Registration was performed to correct for sample drift during acquisition. Scale bar: 10 μm; imaging interval: 10 seconds.

##### Supplemental Movie 2

**Example showing dynamic changes in Galvanin-GFP fluorescence distribution and protrusion/retraction activity.** Phase microscopy, segmentation mask, Galvanin-GFP fluorescence intensity, and protrusion/retraction activity are shown for a HL-60 neutrophil cell expressing Galvanin-GFP. Cells were confined by a 2.5% agarose gell and Imaged using an oil 60x 1.4 NA plan apo phase contrast objective. Cells exposed to an electric field (300 mV/mm) between 5 minutes and 10 minutes. The colors match those shown in Figure 3A. Scale bar: 10 μm; imaging interval: 5 seconds.

##### Supplemental Data Table 1

**CRISPRi screen data table.** The .csv file contains the normalized $\log_2$ fold-changes in sgRNA abundance for the second round of CRISPRi screens of galvanotaxis and undirected cell migration. The sign for each gene indicates whether knockdown led to an enrichment of the associated sgRNA (positive) or depletion (negative), with adjusted p-values calculated as described in Methods.
